# Mining hidden knowledge: Embedding models of cause-effect relationships curated from the biomedical literature

**DOI:** 10.1101/2021.10.07.463598

**Authors:** Andreas Krämer, Jeff Green, Jean-Noël Billaud, Nicoleta Andreea Pasare, Martin Jones, Stuart Tugendreich

## Abstract

We explore the use of literature-curated signed causal gene expression and gene-function relationships to construct unsupervised embeddings of genes, biological functions, and diseases. Our goal is to prioritize and predict activating and inhibiting functional associations of genes, and to discover hidden relationships between functions. As an application, we are particularly interested in the automatic construction of networks that capture relevant biology in a given disease context.

We evaluated several unsupervised gene embedding models leveraging literature-curated signed causal gene expression findings. Using linear regression, it is shown that, based on these gene embeddings, gene-function relationships can be predicted with about 95% precision for the highest scoring genes. Function embedding vectors, derived from parameters of the linear regression model, allow to infer relationships between different functions or diseases. We show for several diseases that gene and function embeddings can be used to recover key drivers of pathogenesis, as well as underlying cellular and physiological processes. These results are presented as disease-centric networks of genes and functions. To illustrate the applicability of the computed gene and function embeddings to other machine learning tasks we expanded the embedding approach to drug molecules, and used a simple neural network to predict drug-disease associations.

## Introduction

Many experimental observations reported in the biomedical literature represent cause-effect relationships. Examples are observations that directly or indirectly couple the activation or inhibition of genes to the downstream regulation of other genes, or the activation or inhibition of biological functions. Collectively, such literature-derived causal relationships (1) can be viewed as the defining features of genes and functions, and therefore be exploited in machine learning (ML) models. A widely used approach is the construction of mappings to high-dimensional vector representations (2), so-called embeddings, that are at the heart of many modern ML methods. The most famous example for this arguably is the word2vec algorithm (3), which uses word proximity in a text to encode semantic relationships in high-dimensional word embeddings. Embeddings have also been applied to graphs (4, 5) and used in scientific contexts, for instance to discover latent knowledge in materials science (6). In the biological context, embeddings for genes have been constructed from protein sequences (7), protein-protein interaction networks (8), co-expression data (9), and using text mining (10, 11).

In this work we explore the use of literature-curated signed causal gene expression and gene-function relationships to construct unsupervised embeddings of genes and functions. In contrast to protein-protein interactions or correlation measures like co-expression, causal gene expression relationships capture information about the behavior of a biological system as a whole in response to perturbations. Here, we make explicit use of the fact that causal interactions carry a sign which distinguishes between activating and inhibiting effects. The obtained gene embeddings can be used to predict and prioritize genes affecting functions and diseases. We distinguish our approach from existing function prediction methods that aim to annotate previously uncharacterized genes with their predicted function, based on some form of “guilt-by-association”, i.e. the assumption that co-localized and interacting genes or proteins are more likely to be functionally correlated (12). Here, in contrast, we are interested in the identification of the most relevant genes causally affecting a given function or disease. These genes can either be previously known to be associated with that function or purely predicted. In the context of diseases, gene prioritization approaches were previously developed based on matrix factorization (13, 14), but those do not distinguish between activating and inhibiting effects. In addition to gene embeddings, we also construct function embedding vectors that allow to infer previously unknown signed function-function relationships, including disease-function associations that point to disease mechanisms and involved cell types or tissues.

Our embeddings are generally useful to construct biological networks that highlight some mechanism or key contexts. A recent example is the “Coronavirus Network Explorer” (15) which uses an early version of our gene-function prediction approach to compute networks that connect SARS-CoV-2 viral proteins to host cell functions. In this paper, we illustrate the application to biological networks by constructing disease networks which capture disease-underlying functions and associated key genes. Embeddings are not limited to genes, but can also be extended to other molecules including drugs. Such embedding feature vectors can then be used in other ML models trained for arbitrary prediction tasks. As an example we demonstrate this for the prediction of drug-disease associations.

## Methods

### Literature-curated content

We employ the QIAGEN Knowledge Base (QKB), a structured collection of biomedical content that includes findings manually curated from the literature as well as content from thirdparty databases (https://digitalinsights.qiagen.com/products-overview/qiagen-knowledge-base/). The QKB was used to create a large-scale knowledge graph with nodes representing genes, chemical compounds, drugs, microRNAs, biological functions, and diseases; and edges categorized into different edge types representing a variety of interactions such as gene expression, activation/inhibition, phosphorylation, and protein-protein binding among others. In this work we particularly use two kinds of edges: 1) gene expression relationships that represent the causal effect of genes on the expression of other genes, and 2) causal gene-function and gene-disease edges that represent causal effects of genes on biological functions and diseases. We only consider edges that have an associated direction of effect which is either activation (leading to an increase) or inhibition (leading to a decrease). All edges generally bundle a number of underlying literature findings from various experimental contexts, therefore edge signs reflect a consensus among all those contexts. As part of an ontology, functions are organized in a hierarchy where, except for very general terms, parents inherit causal gene associations (and edge signs) from their descendants. In total, 6,757 genes and 29,553 functions are included in our embedding model (see Supplementary data, Section 1). Here and in the following, the term “function” generally referes to both functions and diseases, unless we want to explicitly make the distinction.

### Unsupervised gene embeddings

In the following we describe three approaches to derive unsupervised gene embeddings from downstream expression signatures using literature-curated signed causal gene expression relationships. The starting point is a bipartite graph *G* (see Figure 1a) in which *N* genes (for which we will compute embeddings) are connected to their *M* expression-regulated target genes by signed edges that represent causal expression findings from the literature. From *G* we define the signed, weighted *N* ×*M* bi-adjacency matrix *W*, 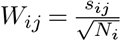, where *s*_*ij*_ ∈ {−1, 0, 1} (activation: +1, inhibition: −1, no edge: 0), and *N*_*i*_ = Σ_*j*_ |*s*_*ij*_| is the total number of genes that are regulated by gene *i*. The matrix *W* can be viewed as taking *N*-dimensional one-hot encoded gene vectors as input and outputting normalized *M*-dimensional vectors corresponding to the up/down regulation pattern (see Figure 1b). Two of our embedding strategies (E1 and E2) are based on an approximation of the matrix *W*, which is associated with the compression of the one-hot encoded input into a lower dimensional embedding space.

**Fig. 1.**
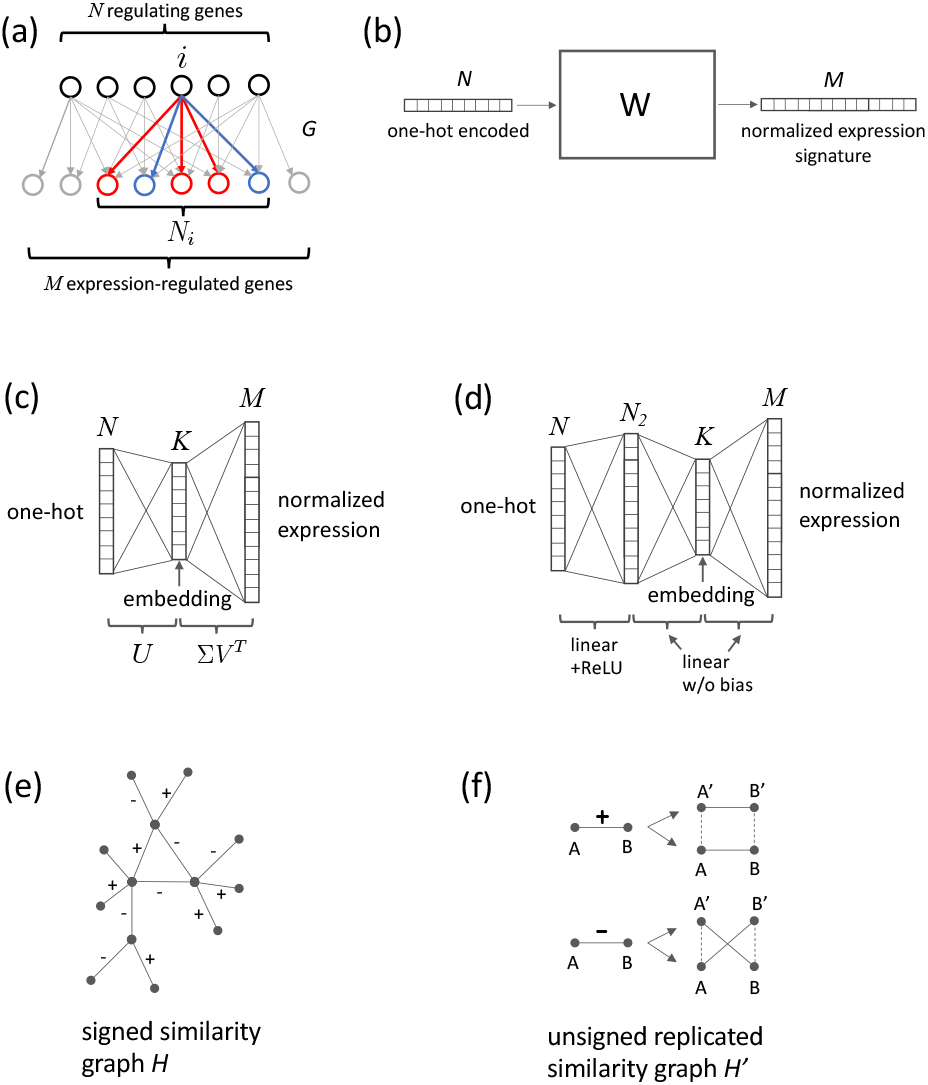
Gene embedding methods. (a) In the bipartite graph *G*, regulating genes are connected to expression-regulated genes by signed edges that represent upregulating (red: +1) and down-regulating (blue: −1) causal expression findings from the literature. Embedding vectors are computed for the *N* regulating genes. *G* defines the signed, weighted adjacency matrix *W*. (b) *W* can be viewed as taking *N*-dimensional one-hot encoded gene vectors as input and outputting normalized *M*-dimensional vectors corresponding to the up/down regulation pattern. (c) The spectral method E1 uses a low-rank approximation 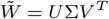 to compute embedding vectors, which is equivalent to training a simple 3-layer linear neural network without bias terms and mean-squared error (MSE) loss. (d) The neural network-based embedding strategy E2 extends the linear model by adding another layer which includes bias and a ReLU activation function. (e) The graph-based approach E3 uses a signed similarity graph *H* connecting similar and anti-similar genes. (f) From *H* an unsigned graph *H*′ is constructed with a replicated set of nodes. *H*′ allows the computation of embeddings using the node2vec algorithm (5).

The “spectral” embedding E1 uses a low-rank approximation of *W* based on singular value decomposition (16),

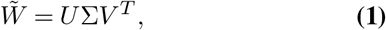

where columns of the *N* × *K* matrix *U* are eigenvectors of the positive definite matrix *S* = *WW*^*T*^, corresponding to its top *K* eigenvalues. Entries of the matrix *S* represent a signed “similarity” of genes based on their downstream regulation patterns. Note, that the normalization factor 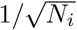 used in the construction of *W* was chosen such that diagonal elements of *S* are equal to one, regardless of the number of regulated genes. The square roots of the eigenvalues of *S* form the matrix elements of the diagonal *K* × *K* matrix Σ, and *V* is a *M* × *K* matrix. One can think of *U* as projecting one-hot encoded vectors representing single genes onto *K*-dimensional embedding vectors, i.e. these embedding vectors are the rows of *U*, where *U*^*T*^ *U* = *I*. This spectral method of computing embedding vectors is equivalent (up to constant scale factors on embedding vector components) to training a simple 3-layer linear neural network without bias terms and mean-squared error (MSE) loss (corresponding to the Frobenius norm of 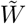), where embeddings are retrieved from the middle layer (17) (see Figure 1c). The neural network-based embedding strategy E2 extends this linear model by adding another layer which includes bias and has a rectified linear unit (ReLU) activation function in order to capture non-linear effects (see Figure 1d). Since there is no bias term between the final layers for both the E1 and E2 approaches, inverting the sign of an embedding vector will result in exactly the opposite effect on downstream regulated genes.

For the third embedding strategy (E3), instead of using the signed similarity matrix *S*, we construct a signed similarity graph *H* that has a signed edge between two gene nodes *i* and *k* if the two genes exhibit a similar downstream regulation pattern. In particular, we compute the “z-score” 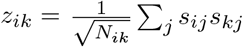 where *N*_*ik*_ = Σ_*j*_ |*s*_*ij*_||*s*_*kj*_| is the number of co-regulated genes, and require the absolute value of *z*_*ik*_ to meet a certain cut off for an edge to be present. The sign of an edge is given by the sign of *z*_*ij*_ (see Figure 1e). From *H* we construct an *unsigned* graph *H*^′^ by replicating each node of *H* and connecting the replicated nodes in *H*^′^ either parallel (positive edge sign) or crosswise (negative edge sign) with unsigned edges as shown in Figure 1f. This construction of an unsigned graph *H*^′^ preserves the information contained in the edge signs of *H*. In the next step we apply the node2vec graph embedding algorithm (5) that samples random walks in order to map the graph embedding problem to word2vec using the skip-gram approach (3). Embedding vectors *u*_*i*_ and *v*_*i*_ are computed for all nodes in *H*^′^, where *u* and *v* denote the two replicas. The final gene embedding vectors are then obtained by taking the difference, *u*_*i*_ − *v*_*i*_ which preserves the same symmetry w.r.t. sign changes as described for the spectral and neural network-based approaches, i.e. a gene with the opposite effect on expression regulation would have an embedding vector whose sign is inverted.

### Function embeddings

Functions are characterized by their causally-associated genes that were curated from literature along with the respective direction of the effect (activation or inhibition). We construct function embedding vectors *p* in the same vector space as gene embedding vectors *x* such that their scalar product *p* · *x* approximates the effect of *x* on *p*, (activation: *p* · *x >* 0, inhibition: *p* · *x <* 0, no effect: *p* · *x* ≈ 0). This construction is in line with the symmetry described above: a gene with opposite causal expression signature, i.e. with the embedding vector −*x* has also the opposite effect −*p* · *x* on the function *p*.

Function embedding vectors are determined as follows: Let the matrix *Y* = {*Y*_*ij*_} represent the effect of gene *i* on a function *j* (activation: *Y*_*ij*_ = 1, inhibition: *Y*_*ij*_ = −1, no effect: *Y*_*ij*_ = 0) as curated from the literature, then the embedding vector *p*_*j*_ for function *j* is determined by standard linear regression (using MSE loss), i.e. minimizing Σ_*i*_ (*x*_*i*_ · *p*_*i*_ − *Y*_*ij*_)^2^. This leads to

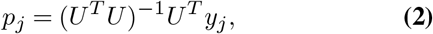

where the matrix *U* has *K*-dimensional gene embedding vectors as rows, *y*_*j*_ is a column vector of *Y*, and it is assumed that the r.h.s. of Equation (2) is well-behaved, and no further regularization is needed, which is usually the case if *K* ≪ *N*. For the spectral method E1 in particular we have *U*^*T*^ *U* = *I*, which simplifies Equation (2) to *p*_*j*_ = *U*^*T*^ *y*_*j*_. Note, that gene-function prediction is viewed as a regression problem, not classification, since the values of *Y*_*ij*_ are ordered in a sequence, −1, 0, 1, and there could in principle be a continuous transition from “inhibition”, to “no effect”, to “activation”. We finalize the construction of function embedding vectors by also performing a normalization step, 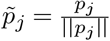, in order to put embedding vectors on the same footing for all functions. This is motivated by the expectation that isotropically distributed random gene embeddings (i.e. “noise”) should lead to the same distribution of 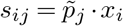 for all functions.

### Gene-function prediction and prioritization

Signed causal gene-function relationships are predicted if the absolute value of the gene-function score defined by the scalar product 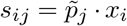 is greater than a certain threshold. For a given function, we can think of function embedding vectors 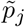, based on the construction above, to be tilted towards “consensus” sets of function-associated genes that have similar (or anti-similar) gene embedding vectors. This means that predicted genes that are also similar to one of these sets, as well as all genes within these sets (that are already known to be associated with the function), will receive high absolute scores. In this sense scoring will prioritize “key” genes that are concordant with the consensus sets. Likewise, genes whose embedding vectors are more scattered and not similar to one of the consensus sets, will not receive high scores, and thus not be prioritized. The choice of the embedding dimension *K* determines whether the gene-function prediction model tends to under- or overfit. If *K* is too small, not enough information will be encoded in the embedding vectors; if *K* is too large, similarity between genes will not be sufficiently represented. For example in the spectral model E1, in the limit *K* = *N* all gene embedding vectors are orthogonal.

Gene-function scores were also transformed to z-scores (see Supplementary data Section 4). Since z-scores measure statistical significance, this is useful to define meaningful cut offs for top-scoring genes.

### Cosine similarity for embedded functions

Similarity of functions is determined by using cosine similarity of the associated embedding vectors, which in our case is simply given by their scalar product since function embedding vectors are normalized. This scalar product can assume negative values corresponding to “anti”-similarity, i.e. the activation of one function being similar to the inhibition of another. Statistical significance of function similarity can be assessed by considering the standard deviation *σ*_*c*_ of the cosine similarity distribution (centered around 0) for two random unit vectors. Since one of these vectors can be held fixed, this is the same as the standard deviation of a single vector component *x*_*i*_ of a random unit vector. From the condition 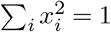 then follows that 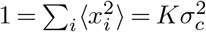 since all *K* vector components are equivalent. An appropriate significance threshold (at 2*σ*_*c*_) for the cosine similarity score is therefore 2*K*^−1*/*2^ which is about 0.09 for a typical embedding dimension of *K* = 500.

### Implementation

Algorithms were implemented in Python using the standard scientific computing stack (numpy, scipy, pandas, scikit-learn). Most code was run on a standard laptop in minutes to hours time frame. The implementation of the neural network-based embedding strategy E2 uses the pytorch framework, and we ran experiments on a machine with a T4 GPU (about 1 hour per run). For node2vec (E3) we utilized the python implementation provided by Grover and Leskovec (5) based on the gensim library with default parameter settings (random walks with 30 nodes, 100 walks per node, hyperparameters *p* = *q* = 1).

## Results

### Cross validation of gene-function prediction

We used the following cross-validation approach to test accuracy of gene-function prediction. We randomly set gene-function relationships *Y*_*ij*_ to zero, trained the linear regression model, and then determined how well those removed gene-function relationships could be predicted. To avoid artificial dependencies between functions we included only “leaves” of the function hierarchy in the subset of functions on which the model was tested, and required that functions were supported by at least 10 genes. A balanced test set was created by randomly picking *n* entries of the matrix *Y* = {*Y*_*ij*_} that had the value 1, *n* entries that had the value −1, and 2*n* entries that were zero. We repeated the procedure *k* times to create *k* in-dependent test sets. For each test set, the selected elements of *Y* were set to zero, and a model was trained using this new matrix *Y*. From the resulting gene-function scores, we then computed receiver-operating characteristic (ROC), and precision-recall curves (PRC). Strictly speaking, zero-entries of *Y*, i.e. the lack of a gene-function relationship in the curated content are not true negative examples in a training or test set, since they do not mean that there was experimental evidence of no functional effect. However, we can assume that the vast majority of zero-entries in *Y* are true negative examples, and the few “false” negative examples do not significantly affect test results.

Two prediction tasks were considered. For the first task, we predicted the presence of a gene-function relationship using an absolute gene-function score threshold |*s*| for the complete test set with 4n examples. For the second task, we used the signed score itself to predict the sign of the effect, i.e. whether it is is activating or inhibiting, and the test set was limited to the 2*n* non-zero examples. There are two sub-cases corresponding to the prediction of either activation (vs. inhibition) or inhibition (vs. activation) among edges with unknown sign, which means there are two distinct PRCs. The ROC is symmetric w.r.t. these two sub-cases, i.e. the second sub-case can be obtained from the first by transforming true (TPR) and false positive rates (FPR) according to TPR → 1 − TPR, and FPR → 1 − FPR, or simply by “flipping” the ROC curve.

Two metrics are used to assess the capability of our signed gene-function prediction model: The AUC, which measures overall how ranking by score discriminates between true positives and negatives, and the precision in the limit of low recall (here set to 5%) which measures how precise the predictions for the highest-scoring genes are. We use the latter metric because we are particularly interested in the identification of the most relevant, key genes causally affecting a given function or disease. In all cross-validation experiments, we set *n* = 1000 and *k* = 50.

Figure 2a shows average AUC and precision at 5% recall for absolute and sign prediction as a function of the embedding dimension *K* for all models E1, E2, and E3. The neural network model E2 uses a single intermediate layer with *N*_2_ = 1000 nodes, and the z-score cut off for the graph-based model E3 was set to *z* = 1.5. Error bars shown correspond to the measured standard deviation across the *k* replicated runs. We observed that increase of the number of nodes in the intermediate layer, or inserting an additional layer (E2) did not result in significant change, and larger cut off values *z* lead to a decrease of AUC and precision (E3). From Figure 2a one can obtain “optimal” embedding dimensions for which AUC and precision are maximal. Embedding dimensions greater than this optimal dimension will lead to over-fitting, while smaller embedding dimensions result in under-fitting of the model. This can be seen for all three cases, E1, E2, and E3, with slightly different behavior of AUC and precision curves. For the spectral case E1 (absolute prediction), the AUC curve shows a very broad peak with maximum AUC ≈ 0.68, while precision (at 5% recall) has a plateau around 95% for dimensions larger than 500, and drops sharply toward lower embedding dimensions. The behaviors of cases E1 and E2 are very close to each other (for absolute and sign prediction) with the AUC (for absolute prediction) dropping slightly more strongly towards high dimensions for the latter. For E3, performance is also similar except that the AUC is lower for absolute prediction, and the maximum (at AUC=0.637) appears shifted to lower embedding dimensions likely because the model included many fewer genes. Figure 2b shows ROC and precision-recall curves for the near optimal cases *K* = 500 (E1), *K* = 350 (E2), and *K* = 100 (E3). All three models reach an average precision of nearly 95% for absolute prediction and about 90% for sign prediction, while the AUC for sign prediction is about 0.70. For the spectral approach E1 we also evaluated models that require each included gene to have a minimum number of downstream regulated genes in the bipartite graph *G* (see Supplementary data, Section 2).

**Fig. 2.**
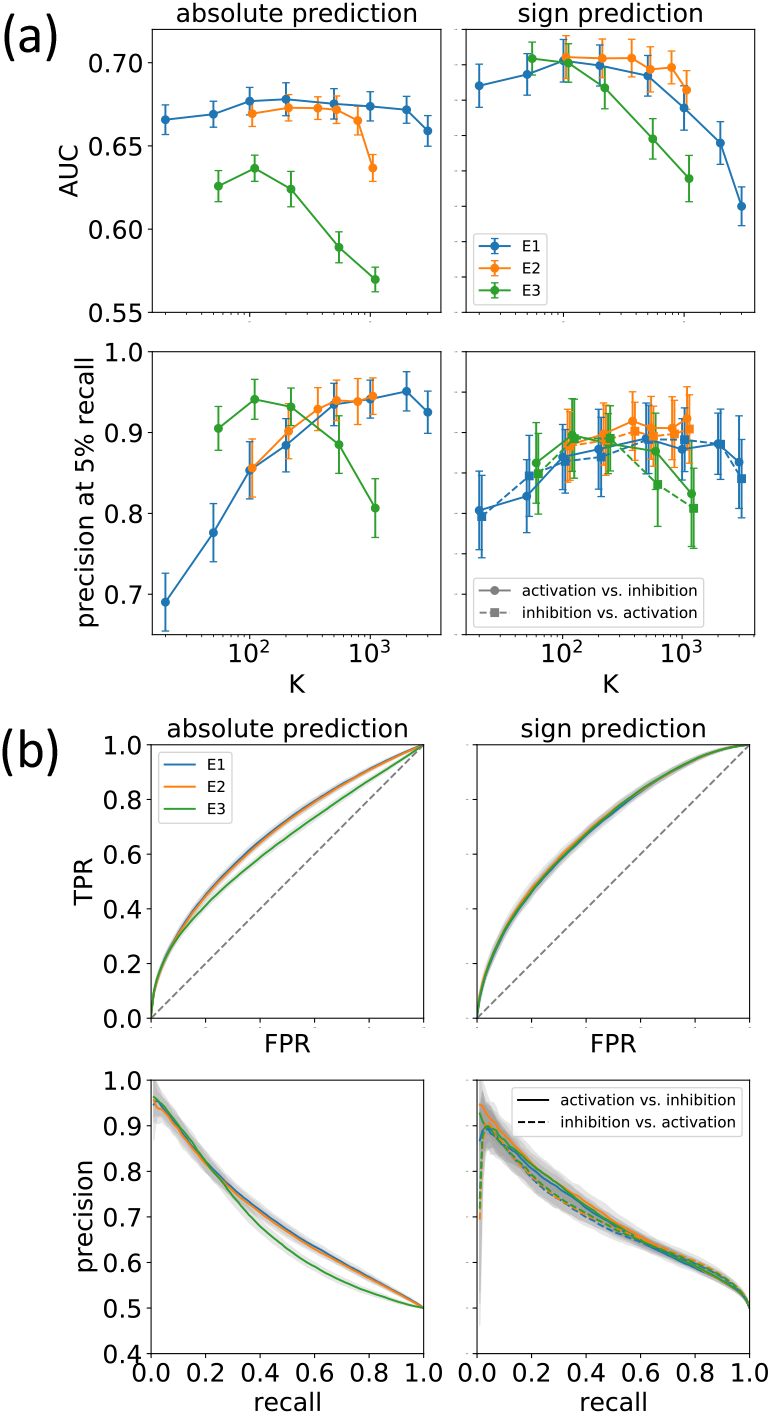
Cross validation: (a) Average AUC and precision at 5% recall for absolute and sign prediction as a function of the embedding dimension *K* for models E1, E2, and E3. (b) ROC and and precision-recall curves for the near-optimal cases *K* = 500 (E1), *K* = 350 (E2), and *K* = 100 (E3). Error bars and shaded areas reflect standard deviations across 50 independent cross-validation runs

As a result we find that there is no advantage of using the computationally more expensive neural network-based (E2) and graph-based (E3) models compared to the spectral model E1. In the remainder of the paper we therefore focus on the spectral model only.

### Function embeddings: discovery of latent biological relationships

Similarity of embedding vectors encoding functions and diseases is expected to reflect underlying biological relationships. In order to test this, we examined how functional contexts are represented in embedding space, constructed a global t-distributed stochastic neighbor embedding (tSNE) map of diseases, and visualized relationships between diseases and associated biological functions (for the latter see Supplementary data, Section 3).

One result of the word2vec algorithm (3) is the association of semantic relationships with simple linear vector operations. For instance, in the most famous example, the vector representation of the word “king” is related to the word “queen” by the (approximate) identity “king” = “queen” − “female” + “male”. In order to find similar relationships in our function embedding space, we consider functions that describe biological processes in a particular context. As an example we examine functions of the form “X of Y”, where the biological process X is from the set *Adhesion, Proliferation, Cell movement, Differentiation*, and Y is a cell type (e.g. *T lymphocytes*, complete list given in Supplementary Table S1). Linear relationships between embeddings can be visualized by performing Principal Component Analysis (PCA), and projecting embedding vectors on the two main principal components which is shown in Figure 3a,b for the process pairs *Adhesion vs. Proliferation*, and *Cell movement vs. Differentiation*. Pairs of functions with different processes, but the same cell type context are connected by straight line segments. If a linear vector relationship like in the “king”-“queen” example above holds, then these line segments are expected to be parallel. From Figure 3a,b it is seen that this is approximately the case for most of the function pairs. In order to make a quantitative assessment of this observation we computed the standard deviation of the distribution of angles that line segments form with the horizontal axis, and compared it to the standard deviation of angles of line segments with randomly shuffled endpoints. The resulting estimated p-values obtained by random sampling are *p* = 1× 10^−5^ for the *Adhesion-Proliferation* pair, and *p* = 4 × 10^−7^ for the *Cell movement-Differentiation* pair, clearly showing the statistical significance of this result.

**Fig. 3.**
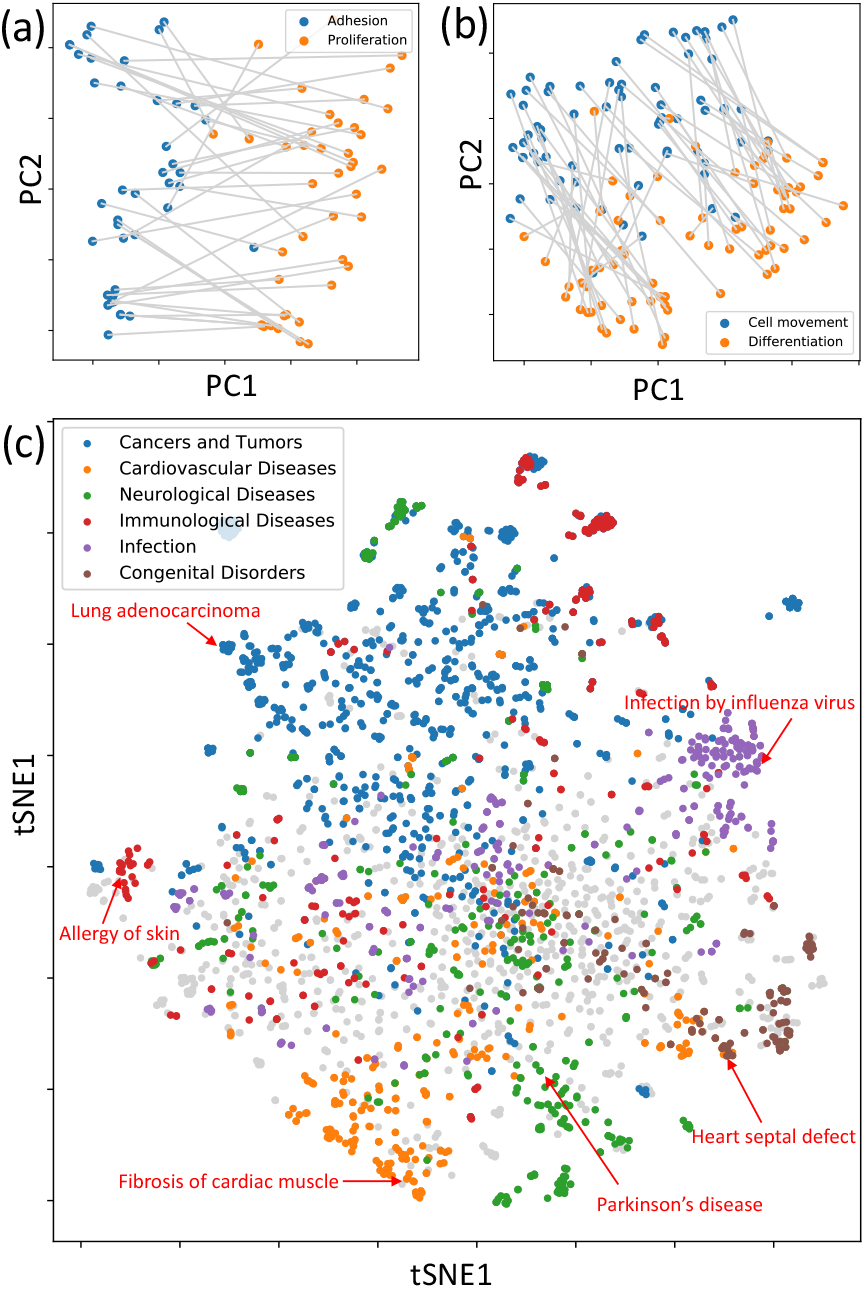
Discovery of latent biological relationships from function embeddings. (a, b) Two-dimensional projection of embedding vectors of functions of the form “X of Y” where X is one of the biological processes *Adhesion, Proliferation, Cell movement*, and *Differentiation*; and Y is one of the cell type contexts given in Supplementary Table S1 (e.g. *T lymphocytes*). (c) Global tSNE visualization of disease embedding vectors. Diseases from different disease categories (cardiovascular, neurological, immunological, infective, congenital, or cancer) tend to cluster together. Note, that cancer and the other disease categories are not exclusive, for instance some cancers were also classified as immunological or neurological, and the non-cancer classification took precedence.

A global tSNE visualization of embedding vectors for diseases (after first reducing dimensionality to 20 using PCA) is shown in Figure 3c. It is seen that, except for the center of the tSNE map, diseases from the same disease category (cardiovascular, neurological, immunological, infective, congenital, and cancer) tend to cluster together, indicating that function embedding vectors capture biological similarity and dissimilarity between diseases.

### Application: inferred disease networks

To explore how the top-scoring genes for a given disease relate to its associated functions, we selected three examples, psoriasis, pulmonary hypertension, and Alzheimer’s disease, which represent a wide spectrum of “systemic” diseases with distinct underlying mechanisms and manifestations. For each of these diseases we determined top-scoring genes and functions and their signs (see tables S2-7 in the supplementary data). In order to give priority to the most “specific” functions (rather than more general terms), we did not include functions that are parents in the process hierarchy of other functions in the list. Redundancy was further decreased by bundling functions from the same context (e.g. cell type), and considering only the highest scoring function from each bundle. For each disease, we constructed a bipartite graph connecting the 15 top-scoring genes and 20 top-scoring functions through edges if the absolute value of the corresponding gene-function score is greater than a certain threshold (here: |z-score|>3), and its sign is consistent with the signs of the adjacent gene and function.

Figure 4 and Figures S4, S5 in the supplementary data show networks constructed this way for all three diseases above. In the following we discuss the psoriasis network. Similar discussions for the other two diseases are given in the supplementary data (Section 4).

**Fig. 4.**
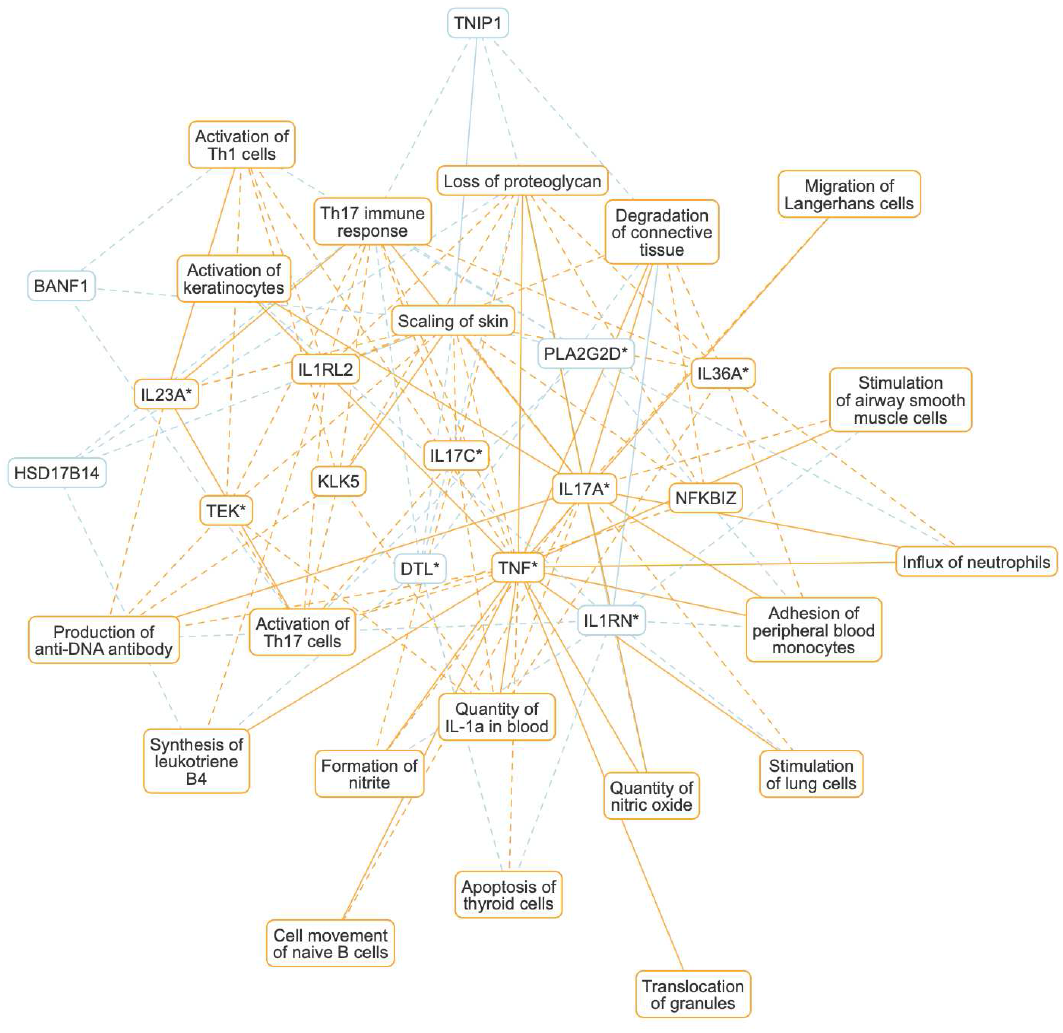
Psoriasis network. Bipartite graph connecting the 15 top-scoring genes, and 20 top-scoring functions through edges with high absolute gene-function scores (|z-score| > 3). The network shows disease-underlying biological functions and known disease genes, as well as genes that are predicted to be implicated in psoriasis based on QKB content. Each node (gene or function) carries a color-coded sign (positive: orange, negative: blue) depending on whether that gene or function is positively- or anti-correlated with psoriasis. The edge style indicates whether gene-function relationships are supported by content of the QKB (solid), or purely inferred (dashed). Genes marked with an asterisk (*) have known associations with psoriasis in the QKB.

Psoriasis is a chronic inflammatory skin disease with a strong genetic component (18). The disease has multiple forms and also may affect organs other than the skin. The network shown in Figure 4 highlights the main immune axis represented by the IL17-IL23 T helper components (*Activation of Th1 cells, Activation of Th17 cells*). IL17 and IL23, as well as TNF, are known to be involved in the pathogenesis of psoriasis. One of the hallmarks of psoriasis is keratinocyte proliferation and immune cell infiltration. This and the disease phenotype (*Scaling of skin, Degradation of connective tissue*) are well represented among the functions shown in the network (*Activation of keratinocytes, Adhesion of peripheral blood monocytes, Cell movement of naive B cells, Influx of neutrophils, Migration of Langerhans cells*). A number of genes shown are purely predicted from QKB content (BANF1, HSD17B14, IL1RL2, KLK5, NFKBIZ, TNIP1). An independent literature search uncovered known or suspected involvement of these genes in the disease: BANF1 has been suggested to be associated with upregulated proliferation of keratinocytes in psoriatic lesions (19). Kallikreins (like KLK5) were found in the serum of patients with psoriasis which suggests that they might be involved in the pathogenesis (20). The expression of NFKBIZ (a nuclear inhibitor of NF-*κ*B) in keratinocytes has been found to trigger not only skin lesions but also systemic inflammation in mouse psoriasis models (21). Loss of TNIP1 in keratinocytes leads to deregulation of IL-17-induced gene expression and increased chemokine production in vitro and psoriasis-like inflammation in vivo (22).

This demonstrates that these networks indeed capture known underlying disease mechanisms and have the potential to generate novel insights.

### Application: drug-disease prediction

In the following, we demonstrate that the embeddings computed with our approach can also be used for independent prediction tasks. As an example, we built a simple ML model, trained on drug-disease pairs collected from drug labels or clinical trial information, to predict drug effects on diseases. Since the QKB also contains literature-derived information about the effect of drugs on gene expression, it is straightforward to extend the gene embedding model to drug molecules by simply including them in the bipartite graph *G* (see Methods section). Using the spectral model (*K* = 500), we constructed embedding vectors for 1077 drugs and 1388 diseases that were included in the QKB from curated drug labels or clinical trials. These embeddings were combined to build “compound” feature vectors for arbitrary drug-disease pairs (see Figure 5a). No drug information was used for the disease embedding vectors, which were constructed using the standard spectral approach outlined in the previous sections. Using the compound feature vectors, we then trained a multilayer perceptron (MLP) to predict novel drug-disease associations. Two different training sets were considered: one containing 2,102 drug-disease pairs curated from drug labels only, and one also including drug-disease pairs from clinical trials (13,182 drug-disease pairs in total). The same number of negative training samples was randomly drawn from the set of all possible drug-disease combinations. For both training sets we performed cross-validation on 10% of the drug-disease pairs randomly held out and repeated the experiment 50 times. We also considered the (harder) task of predicting drugs or diseases not seen in the training set before, by randomly excluding 50 drugs (or diseases) from the training set, and predicting drug-disease pairs only on those. Resulting average ROC and precision-recall curves are shown in Figure 5b and c: For the training set based on drug labels only and random hold-out, the AUC is 0.857 (0.872 when clinical trials are included), restricting this to only cancer-related diseases in the test set increases the AUC to 0.922 (drug labels only). For the harder prediction task involving only drugs or diseases unseen during training these values are significantly lower, for new drugs AUC=0.725, for new diseases AUC=0.788 (drug labels only).

**Fig. 5.**
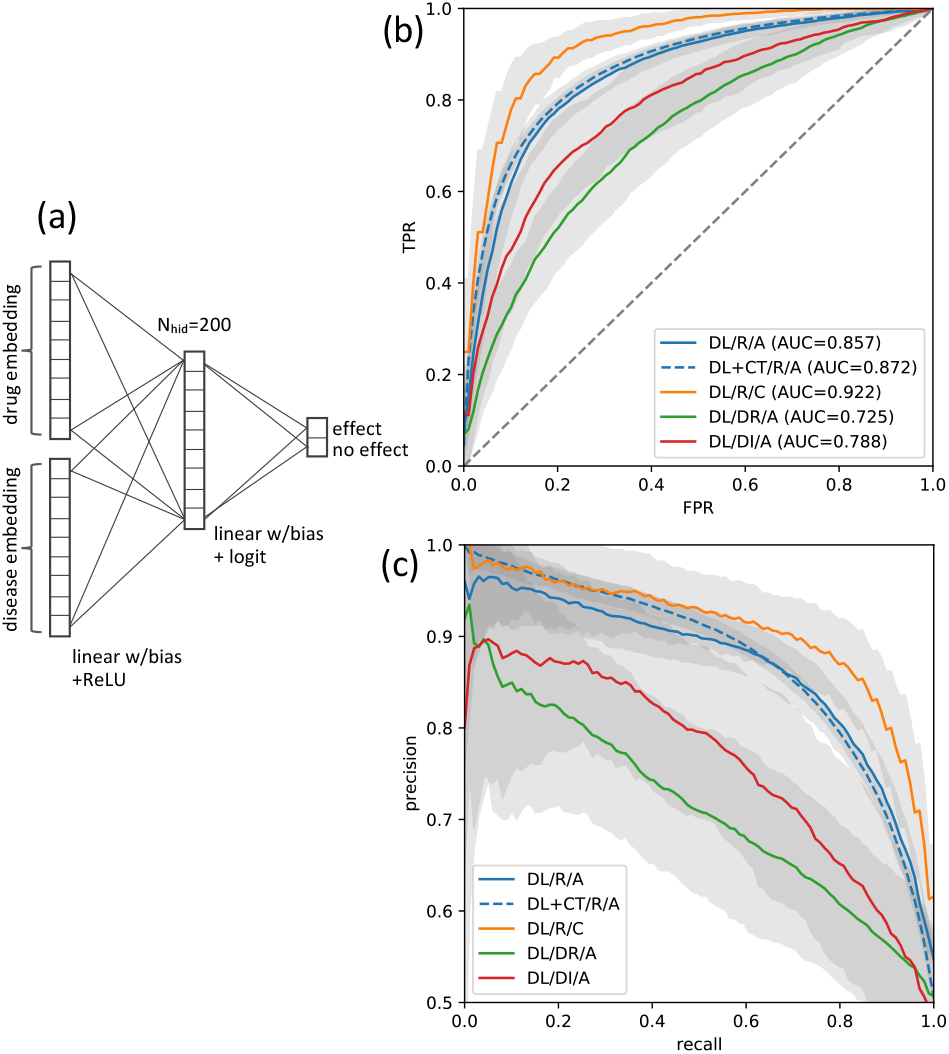
Drug-disease prediction. (a) MLP trained on drug labels and/or clinical trial information to predict drug-disease associations using feature representations based on both, drug and disease embedding vectors. The MLP used here has one hidden layer with 200 nodes. The following cases were considered (see legends): random pairs in test set (R), only drugs included in the test set that were not seen during training (DR), and only diseases included in the test set that were not seen during training (DI). We also distinguished between training based on drug labels (DL) only and drug labels plus clinical trials (CT), as well as all diseases (A) and cancer-related only (C). (b) Average receiver operating characteristics. (c) Average precision-recall curves. Shaded areas reflect standard deviations across 50 independent cross-validation runs.

### Comparison to gene embeddings based on other information

We compared our gene embeddings to those obtained with gene2vec (9) (based on co-expression) and Mashup (8) (based on protein-protein interactions). For the gene-function prediction task (see Results section) we find that our approach outperforms gene2vec, while performing at the same level as Mashup. We also find that top scoring gene sets computed with our approach are mostly disjoint from those computed with Mashup. For a discussion see Supplementary data, Section 5.

## Discussion

We have used signed cause-effect relationships curated from the biomedical literature to construct high-dimensional embeddings of genes, biological functions, and diseases. Gene embeddings are based on literature-derived downstream expression signatures in contrast to embeddings obtained with existing approaches that leverage either co-expression, or protein binding networks. Function embeddings are constructed using gene embedding vectors with a linear model trained on signed gene-function relationships.

Three separate methods were applied to construct gene embeddings, a “spectral” approach based on a low-rank matrix approximation, a neural network-based approach to capture non-linear effects, and a graph-based method utilizing the node2vec algorithm. All three methods performed similarly, reaching on average close to 95% precision for top-scoring genes (90% precision for distinguishing between activating and inhibiting effects) in cross-validation experiments for the gene-function prediction task.

By analyzing various examples, we showed that function embedding vectors capture hidden biological relationships as well as semantic context similar to word embeddings. As an application, we determined top-scoring genes and related functions for three diseases, Alzheimer’s disease, pulmonary hypertension, and psoriasis, to build disease-specific networks. These networks show key genes known to be involved in disease progression, and they capture underlying cellular and physiological processes. We were able to predict a number of disease genes that were not present in the training data (i.e. connected to the disease in the QKB) but could be validated through an independent literature search. It shall be noted that a current constraint of our method is that only a fraction of genes (≈30%) can be covered, limited by content curation and available literature coverage.

In order to demonstrate the applicability of our approach to other prediction tasks, we extended gene embeddings also to drug molecules, and used a simple MLP, trained on known drug-disease associations from drug labels and clinical trials, to predict new drug-disease associations. In cross-validation testing using random drug-disease pairings as negative training set, here, we achieved an average AUC of 0.872.

Our work illustrates that prior knowledge from the biomedical literature can be used collectively to generate new insights, going beyond the findings reported in individual research articles. Applications of knowledge-driven embedding models are manifold. As already implied by the disease networks discussed here, the approach can be used to create new hypotheses for biological mechanisms, identify new potential gene targets for drug repurposing, or predict possible new disease indications in a given therapeutic context.

## Acknowledgments

We would like to thank Dan Shiffman, Jamie Hill, and Paula Tataru for guidance and discussion.

## Supplementary Data

### 1. Knowledge graph summary statistics

The knowledge graph obtained from the QKB and used for this paper contains in total 6,757 genes that appear in both literature-derived gene expression and gene-function relationships. The total number of included expression edges is 147,792 with 286,022 underlying literature findings, and 14,176 regulated genes. There are 217,239 gene-function edges with 395,224 underlying findings that regulate 29,553 functions of which 7,388 are diseases. As part of an ontology, functions are organized in a hierarchy where, except for very general terms, parents inherit causal gene associations (and edge signs) from their descendants. This inheritance mechanism increases the total number of gene-function edges to 748,626. The sign distribution on edges is slightly unbalanced, with roughly two thirds of edge signs being positive, and one third of edges being negative (for both, gene expression, and gene-function edges).

### 2. Cross-validation for smaller networks

For the spectral model E1 we also built models that require each included gene to have a minimum number *N*_*min*_ of downstream regulated genes in the bipartite graph *G*. The number of embedded genes included in these models decreases strongly with increasing value of *N*_*min*_ which is shown in the inset of Figure S1. Because the spectral model is linear, it is expected that the optimal embedding dimension scales linearly with the number of genes (i.e., the size of the matrix *S*). This is confirmed by overlaying scaled AUC-vs-dimension and precision-vs-dimension functions in one plot showing an approximate collapse onto one curve (see Figure S2). Figure S1 shows AUC and precision (at 5% recall) plotted against the parameter *N*_*min*_ for these optimal embedding dimensions. It is seen that the AUC for absolute prediction decreases for increasing *N*_*min*_ because functions are represented by fewer genes and therefore embedding vectors carry less information. At the same time the AUC for sign prediction increases, presumably because only genes are included that are encoded based on a greater number of downstream expressed genes in *G*, thus reducing noise.

**Figure S1.**
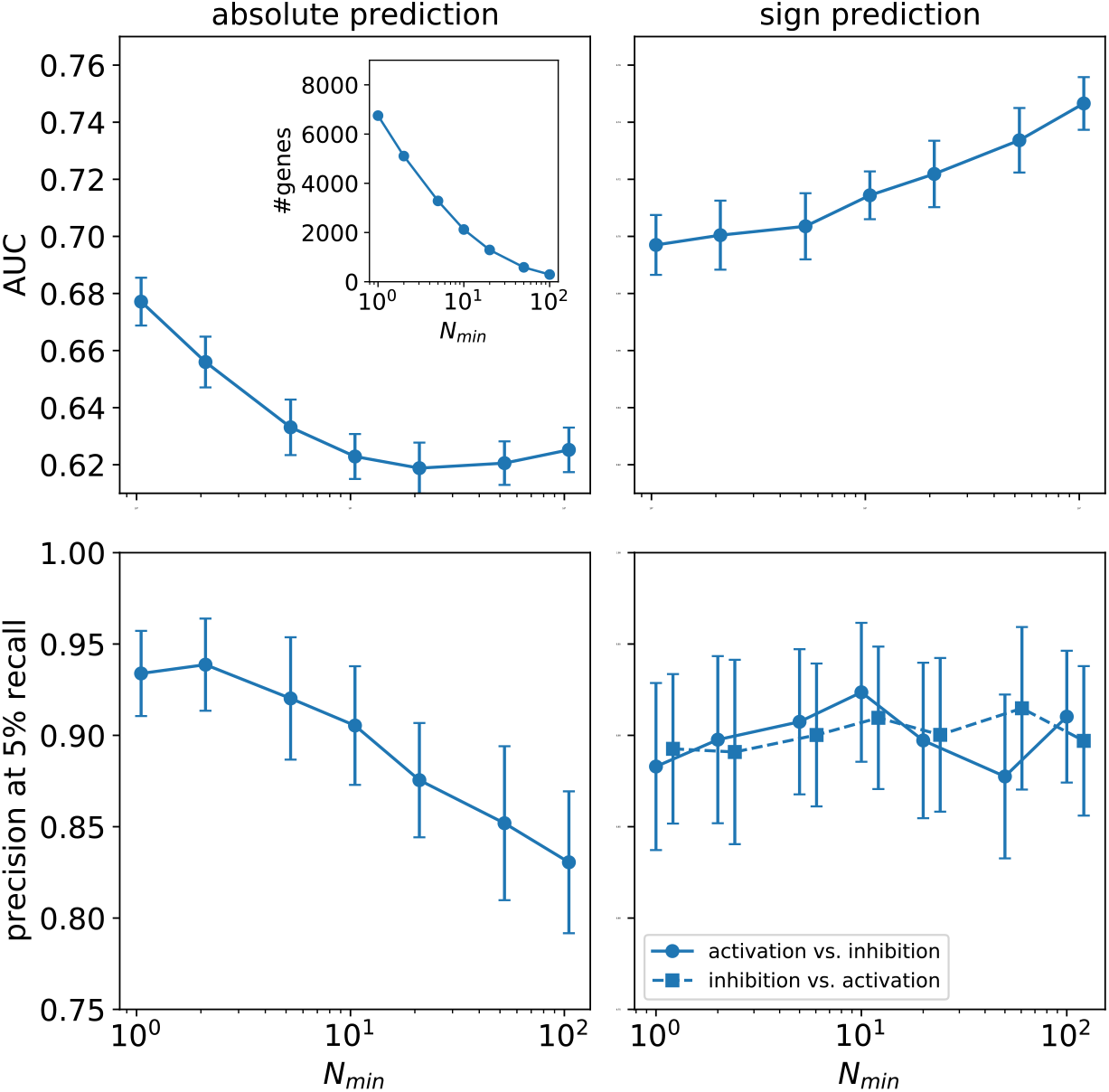
Cross validation: AUC and precision (at 5% recall) plotted against the parameter *N*_*min*_ for optimal embedding dimensions (spectral model). Error bars correspond to the measured standard deviation across the 50 replicated runs. The inset shows the number of genes included in the model as a function of *N*_*min*_.

**Figure S2.**
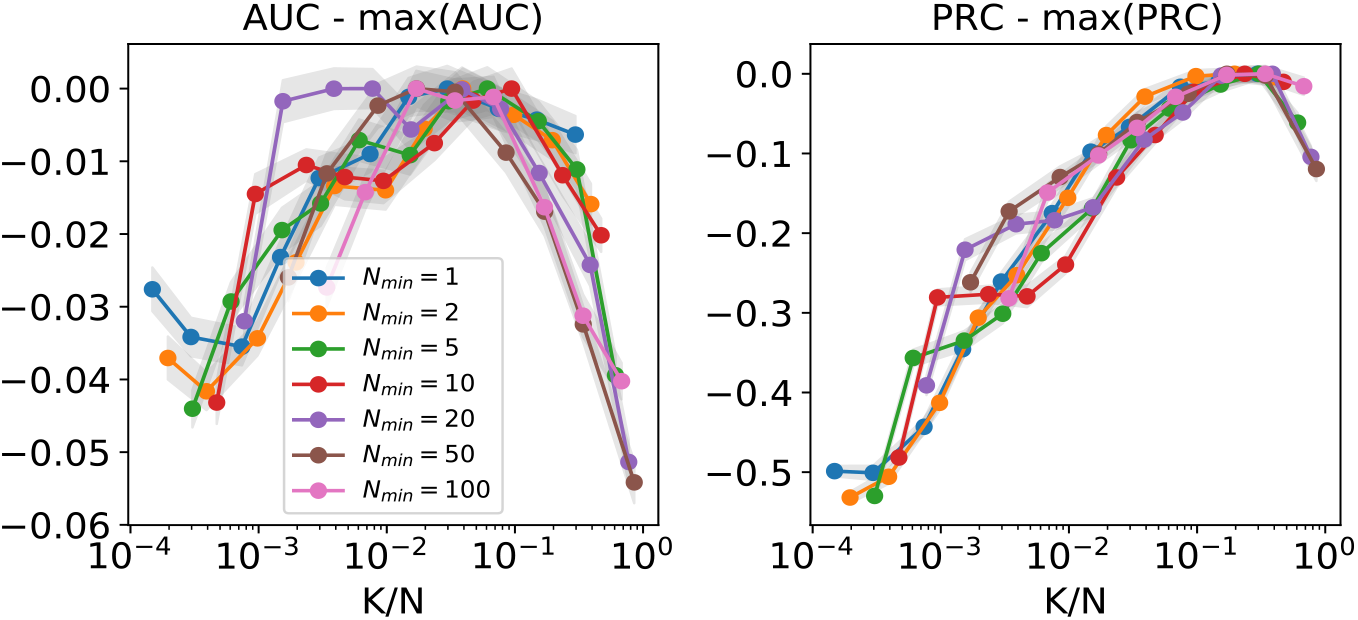
Scaling plots. Because the spectral model is linear, it is expected that the optimal embedding dimension scales linearly with the number of genes. This is confirmed by plotting AUC and precision at 5% recall as function of *K*/*N* (where *K* is the embedding dimension and *N* is the total number of genes) for models with different parameters *N*_*min*_ and also subtracting their maximal value. It is seen that the resulting curves for each model approximately collapse onto one curve.

### 3. Visualization of relationships between a disease and associated biological functions

The neighborhood in embedding space around a given disease can be mapped onto the two top PCA components, where PCA in this case is performed on all functions in the vicinity of the disease, as determined by a preselected cut off on the absolute value of the cosine similarity (here: 0.2). Before performing PCA, we multiply with −1 all function embedding vectors whose cosine similarity is less than zero; those functions *f* are therefore read as “inhibition of *f* “ or “decrease of *f* “. This allows for functions that are related but have an opposite sign to appear close to each other in the PCA projection. The result is shown in Figure S3 for the example of Alzheimer’s disease (AD), where related functions whose sign was inverted are shown in blue, and all others in red. For better readability, Figure S3 shows a subset of all functions, similar to AD, where redundant other functions were removed. This is described in more detail in section 3.3 (main text). Interestingly, this procedure automatically detects many of the underlying disease manifestations of AD which are purely inferred from our embedding model of genes and functions. No explicit function-function or function-disease relationships from the literature were used in this approach. Note, that predicted functions potentially reflect underlying disease mechanisms, however, this cannot be distinguished from processes that share biological aspects of the disease but are not directly involved in it.

**Figure S3.**
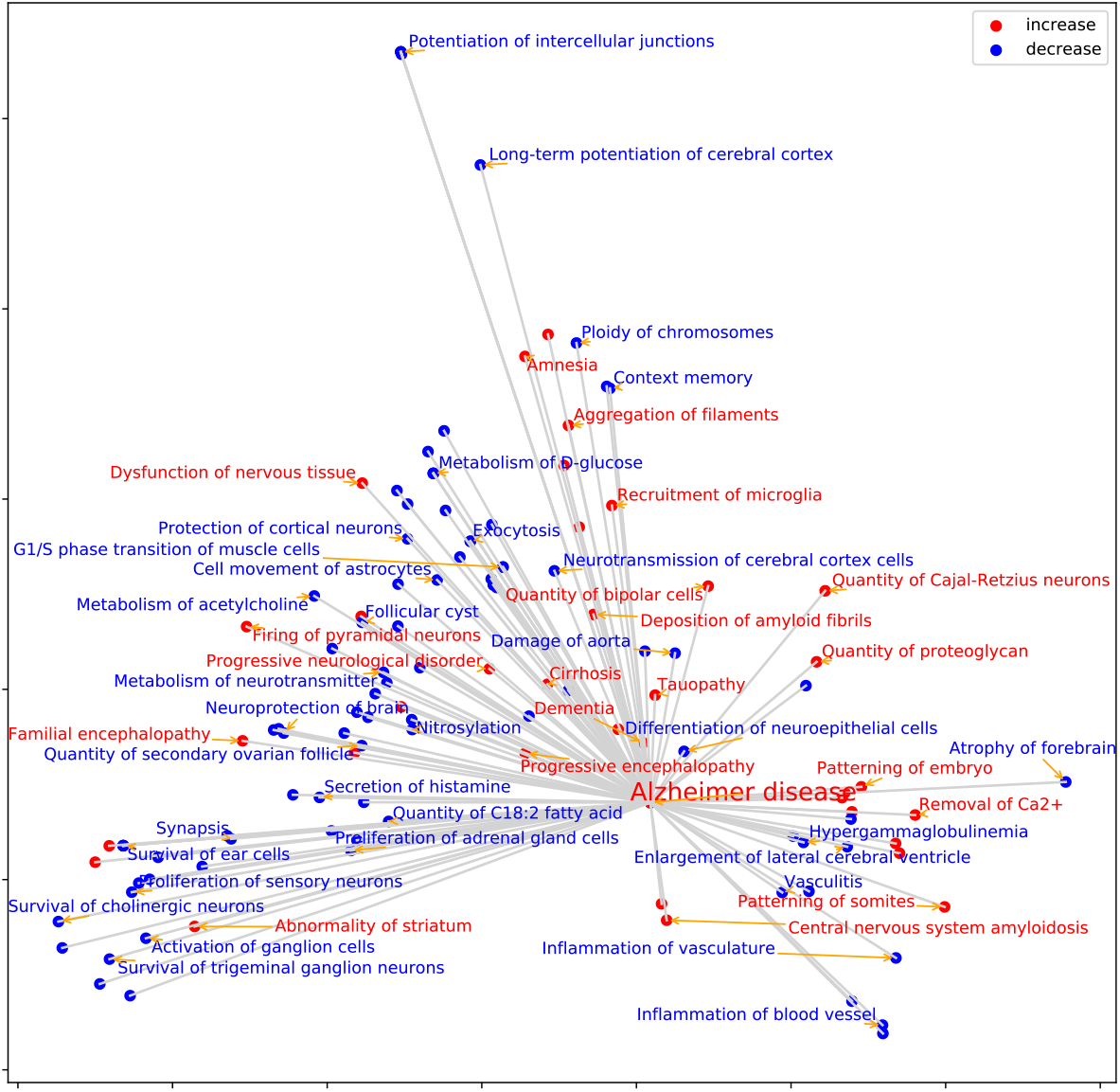
PCA projection of functions and diseases in the neighborhood of “Alzheimer’s disease”. Functions with embedding vectors that are anti-similar to Alzheimer’s disease are shown in blue (with the embedding vector multiplied by −1), vectors that are similar to Alzheimer’s Disease are shown in red.

### 4. Disease networks

For the following, gene-function scores *s*_*ij*_ were transformed to z-scores, 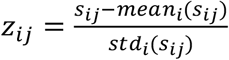 that normalize the distribution of scores *s*_*ij*_ for each function *j* independently. Since z-scores measure statistical significance, this is useful to define meaningful cut offs for top-scoring genes (e.g., |*z*|>2). We verified that z-scores, and the original gene-function scores *s* are linearly related with approximately the same scale factor for all functions, so they can be used interchangeably.

#### 4.1. Alzheimer’s disease

AD is a progressive neurodegenerative disease characterized by severe cognitive impairment, progressive extensive neuronal death, and eventually severe dementia [1, 2, 3]. Neuronal death is one of the main histological and biological markers of AD with hippocampus and striatum being CNS targets, which is reflected in several inhibited functions (i.e., with negative sign) shown in the network in Figure S4 (*Activation of spinal neuron, Cell viability of striatal neurons, Survival of trigeminal ganglion neurons, Chemotaxis of axons, Synaptic transmission of hippocampal neurons, Metabolism of acetylcholine*). Other functions present in the network are *Amyloidosis, Aggregation of filaments*, and *Quantity of proteoglycan* (abnormal amyloid peptide aggregation is a histopathological hallmark of AD), decrease of *Spatial learning* (reflecting cognitive impairment), *Inflammation of vessel* (one of the hallmarks of AD is modification of the cerebral vasculature [4]), inhibition of *Metabolism of D-glucose* (evidence suggests that glucose hypometabolism may be a key player in dementia pathology [5]), and *Acidification of lysosome* (AD is associated with autophagy anomalies, and defective lysosomal acidification contributes to proteolytic failure [6]). The network contains a number of genes that have been implicated in AD and are represented in the QKB (APOE, APP, BDNF, HMGCR, INS, NGF, PSEN1, PSEN2), as well as others that are predicted (CUX2, FBXL7, HRG, LOX, PRR5, SLITRK5, Slfn1). Among the predicted genes, SLITRK5 has no known association with AD per se, but may participate in disease progression through intermediate connections. In fact, SLITRK5 indirectly modulates BDNF, and has a known role in other neurological disorders [7]. Another predicted gene, LOX (lipoxygenase) is known to promote neuroinflammation, and is regarded as a promising therapeutic target for AD [4].

**Figure S4.**
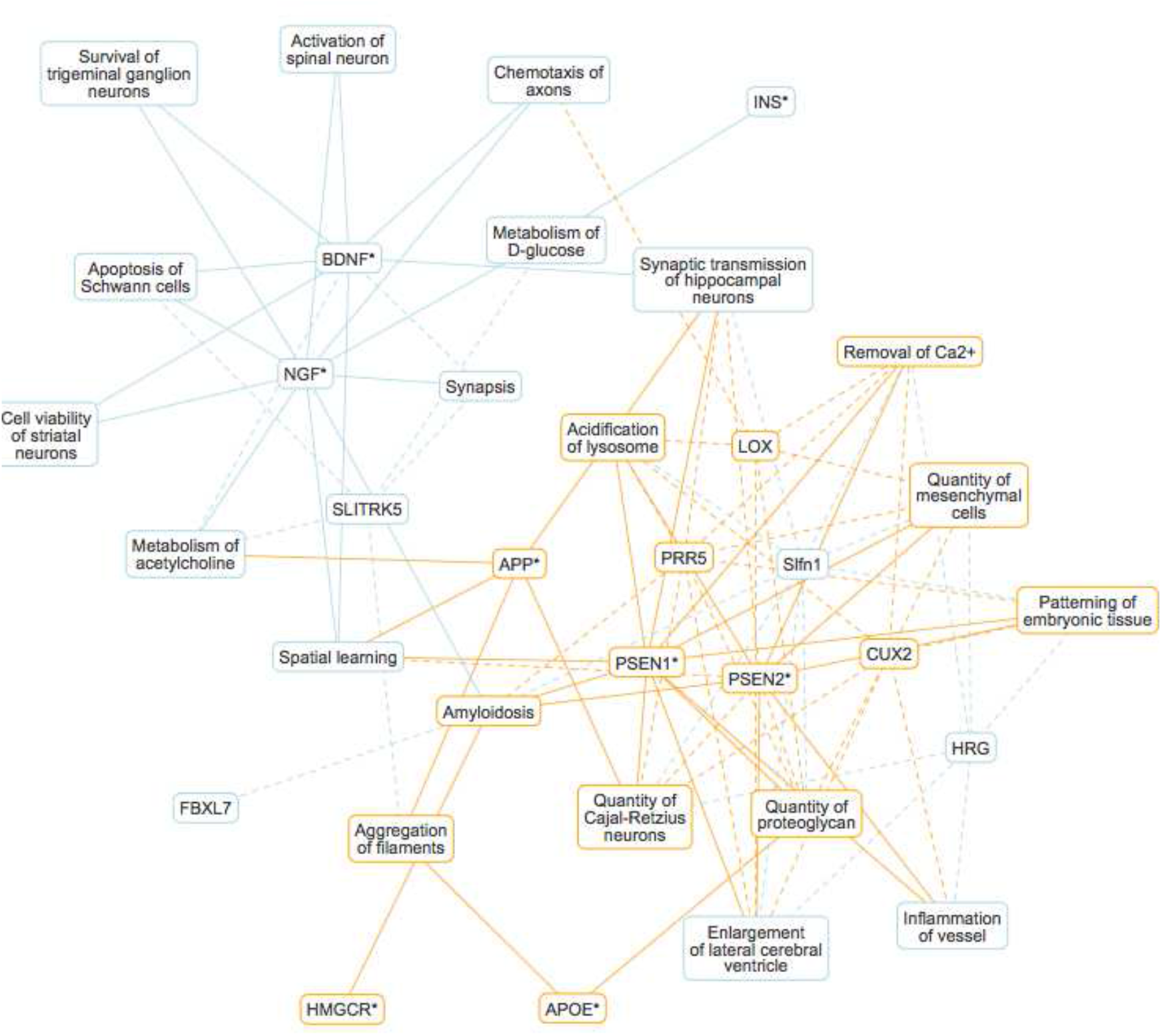
Alzheimer’s disease network. Bipartite graph connecting the 15 top-scoring genes, and 20 top-scoring functions through edges with high absolute gene-function scores (|z-score| > 3). The network shows a number of disease-underlying biological functions and known disease genes, as well as genes that are predicted to be implicated in AD based on QKB content (see detailed discussion in main text). Each node carries a color-coded sign (positive: orange, negative: blue) depending on whether that gene or function is positively- or anti-correlated with Alzheimer’s disease. The edge style indicates whether gene-function relationships are supported by content of the QKB (solid), or purely inferred (dashed). Genes marked with an asterisk (*) have known associations with Alzheimer’s disease in the QKB.

#### 4.2. Pulmonary hypertension

Pulmonary hypertension (PH), especially pulmonary arterial hypertension develops after the resting threshold pressure in pulmonary arteries is exceeded, typically resulting in right ventricular dysfunction and failure, and often leading to death [8]. PH induces vascular remodeling characterized by production of new endothelial cells, myofibroblasts, vascular smooth cells, extracellular matrix changes and fibrosis induction. The network shown in Figure S5 reflects this through appearance of the functions *Systolic pressure of right ventricle, Muscularization of artery*, and *Pulmonary fibrosis*. We also observe roles of immune cells (bone marrow-derived dendritic cells, T lymphocytes, mast cells, macrophages, and others) which are present in vascular lesions in patients with PH [9]. In particular, the recruitment of macrophages in perivascular regions of pulmonary arteries has been observed [10]. A number of genes present in the network have a known association with PH as represented in the QKB (ADA, ADORA2B, APOE, ARG1, BMPR2, IL18, IL1RN, PTGDR2, RPTOR). Among these is ADORA2B, which through its effect on the Pulmonary fibrosis or aplastic anemia appears to mediate the development of PH, and which is regarded as a potential therapeutic target [11, 12]. BMPR2 is also a major player in PH [13], as mutations in the gene have been identified as the main genetic cause [14, 15]. The network indicates the contribution of these two proteins towards increasing the systolic pressure of the right ventricle. The network also contains several predicted genes (ACVR2A, AKNA, AQP11, CCR3, IL15, P4HTM) that are not associated with PH in the QKB. An independent literature search found that aquaporins (AQP0-12) may be involved in PH under hypoxic conditions [16], and ACVR2A is a type 2 BMP receptor like BMPR2.

**Figure S5.**
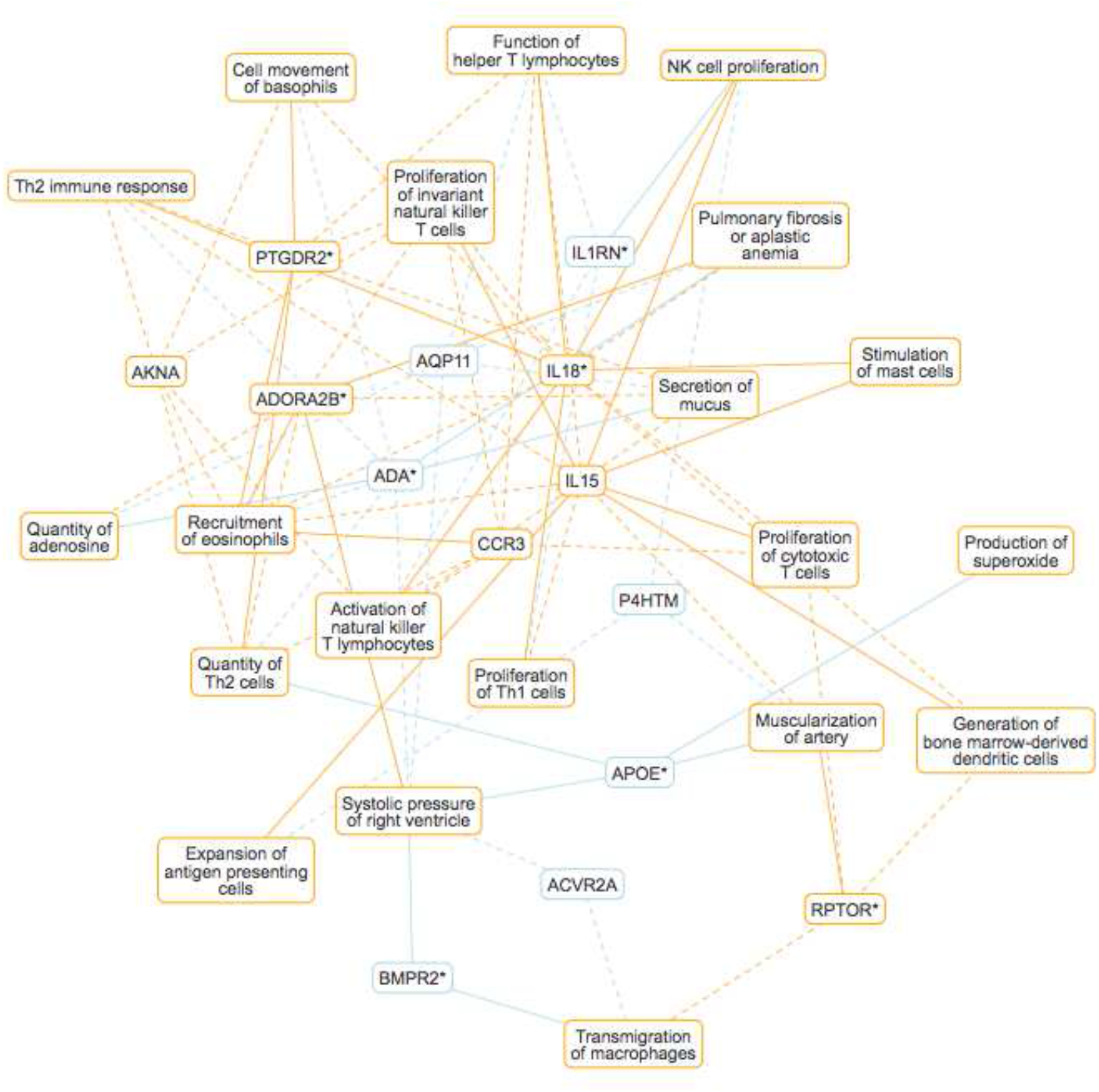
Pulmonary hypertension network. Bipartite graph connecting the 15 top-scoring genes, and 20 top-scoring functions through edges with high absolute gene-function scores (|z-score| > 3). The network shows a number of disease-underlying biological functions and known disease genes, as well as genes that are predicted to be implicated in PH based on QKB content (see detailed discussion in main text). Each node (gene or function) carries a color-coded sign (positive: orange, negative: blue) depending on whether that gene or function is positively- or anti-correlated with pulmonary hypertension. The edge style indicates whether gene-function relationships are supported by content of the QKB (solid), or purely inferred (dashed). Genes marked with an asterisk (*) have known associations with pulmonary hypertension in the QKB.

### 5. Comparison to gene embeddings based on other information

In order to compare our causal expression-based gene embeddings to other gene embedding approaches, we downloaded pre-trained gene embedding vectors generated with both, the gene2vec algorithm (https://github.com/jingcheng-du/Gene2vec), and Mashup (http://cb.csail.mit.edu/cb/mashup). Gene2vec (Du *et al*., 2019) uses gene co-expression patterns from 984 GEO datasets to construct 200-dimesional vector representations of human genes by training a 3-layer neural network with gene pairs that are highly co-expressed. Mashup (Cho *et al*., 2016) is based on a network diffusion approach that performs random walks with restart, and computes a lower-dimensional approximation of diffusion states. Here, we used the pre-computed, 800-dimensional embedding vectors for human genes based on protein-protein interactions from the STRING database [20]. We tested the gene2vec and Mashup embedding vectors on the same gene-function prediction tasks and cross-validated as described in Section 3. These tests were performed on the intersection of genes included in gene2vec (or Mashup) and our spectral model (gene2vec: 6,187, Mashup: 5,689), and for comparison we also reran tests for our model only including genes in the intersection.

For the absolute prediction task gene2vec reaches an AUC of 0.536 and precision at 5% recall of 0.647 compared to values of 0.677 and 0.934 in our model. The superior performance of our model shows that gene encodings based on causal expression responses likely contain more information about gene function than encodings based on co-expression. For Mashup we obtain an AUC of 0.663 and precision at 5% recall of 0.972, i.e. a performance similar to our model (AUC=0.672, precision at 5% recall=0.927). Performance for the sign prediction task was almost identical for both models. This indicates that causal gene expression and protein-protein interactions are on the average equally informative for gene-function prediction when tested on the same set of genes.

To further compare the causal expression-based approach with Mashup (in this case including all 18,362 genes covered by Mashup) we determined the top-scoring genes for all three diseases discussed in Section 3.3 with both methods, focusing only on genes that are predicted, i.e., not already associated with the disease in the QKB. Results are presented as heat map plots in Figure S6, showing that top-scoring gene sets computed for both methods are mostly disjoint. For psoriasis a number of top genes obtained with Mashup (CXCR2, MMP9, FLT4, IL12B) are also picked up (with lower scores) by our approach, however some with the opposite sign. There is also some overlap for pulmonary hypertension where the causal approach also gives high scores for Mashup’s top scoring genes ACVR2A and IL15. In the network discussions in Section 3.3 (main text) and Section 4 (here) we showed that some of our predicted genes could be verified through an independent literature search (AD: SLITRK5, LOX; PH: AQP11, ACVR2A; psoriasis: BANF1, KLK5, TNIP1). Likewise, some genes predicted by Mashup have been associated with the respective disease (AD: GSAP [17], APH1A [18]; PH: TGFPR1 [19]). By and large both approaches appear to be mostly complementary, highlighting the crucial difference in the underlying information used to encode genes, on one side relationships between indirect causal effects on expression signatures, and on the other local molecular interactions. This suggests a possible integration of both approaches in future work.

**Figure S6.**
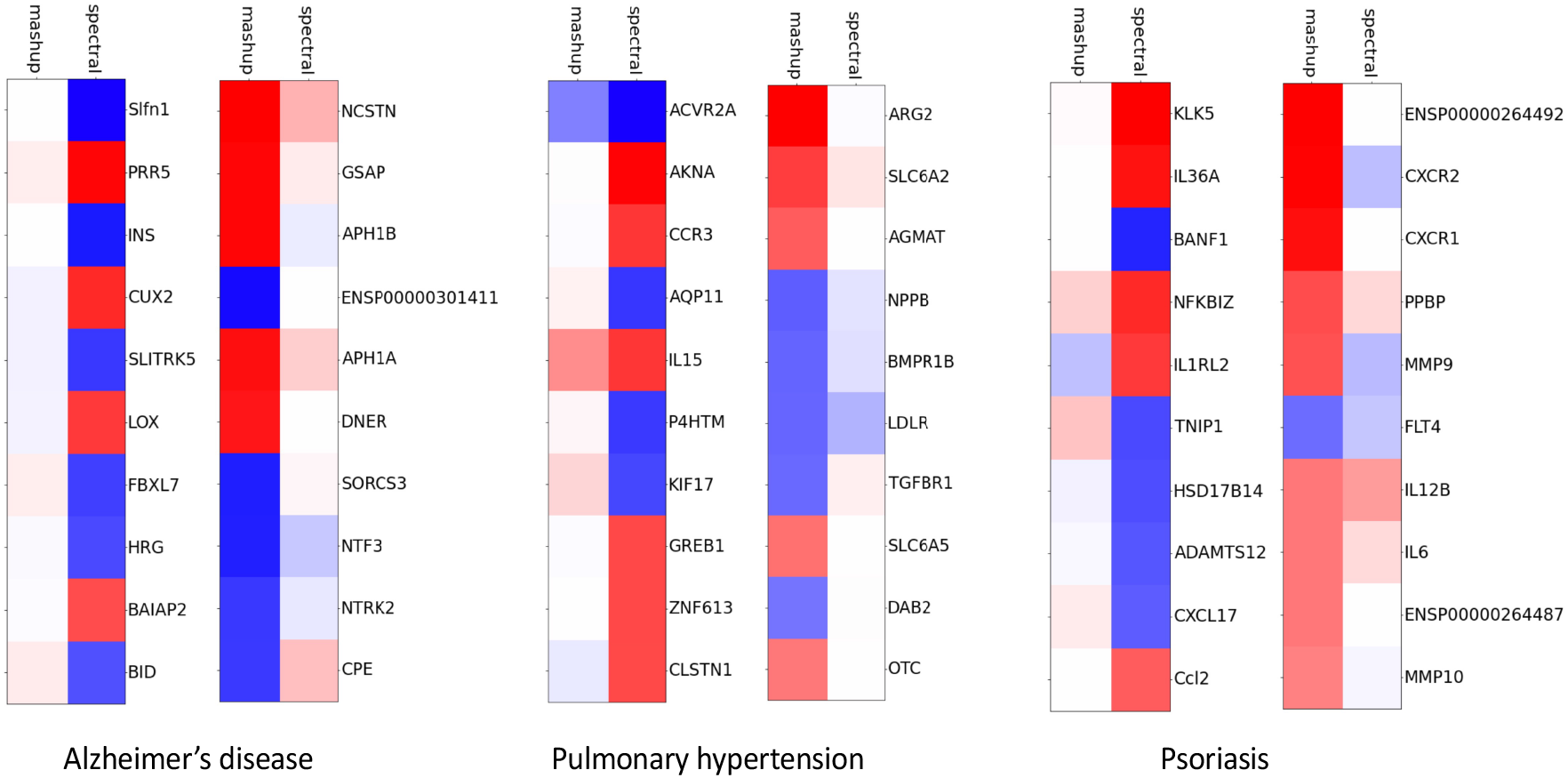
Comparison to Mashup. Top-scoring predicted genes obtained with the spectral model and Mashup for the diseases Alzheimer’s disease, pulmonary hypertension, and psoriasis discussed in Section 3.3. Results are presented as heat map plots showing that top-scoring gene sets computed for both methods are mostly disjoint. For psoriasis, several top genes obtained with Mashup (CXCR2, MMP9, FLT4, IL12B) are also picked up (with lower scores) by our approach, however some with the opposite sign. There is also some overlap for pulmonary hypertension where the causal approach also gives high scores for Mashup’s top scoring genes ACVR2A and IL15. (Red: activated, blue: inhibited)

## Supplementary Tables

**Table S1(a).** Included cell type contexts. Adhesion vs. Proliferation (see Figure 3a)

|  |  |
| --- | --- |
| mononuclear leukocytes | lymphoid cells |
| monocytes | neutrophils |
| dermal cells | lymphocytes |
| granulocytes | myeloid cells |
| PBMCs | leukemia cell lines |
| fibroblast cell lines | bone marrow cell lines |
| immune cells | lymphatic system cells |
| keratinocytes | cancer cells |
| microvascular endothelial cells | endothelial cells |
| peripheral blood leukocytes | smooth muscle cells |
| kidney cells | macrophages |
| epithelial cells | muscle cells |
| embryonic cells | tumor cells |
| mast cells | peripheral blood lymphocytes |
| embryonic cell lines | epithelial cell lines |
| B lymphocytes | lung cancer cell lines |
| gonadal cell lines | phagocytes |
| prostate cancer cell lines |  |
| antigen presenting cells |  |
| red blood cells |  |
| blood cells |  |
| epithelial tissue |  |
| endothelial cell lines |  |
| connective tissue cells |  |
| epidermal cells |  |

**Table S1 (b).** Included cell type contexts. Cell movement vs. Differentiation (see Figure 3b)

|  |  |  |
| --- | --- | --- |
| lymphoma cell lines | B lymphocytes | bone marrow cells |
| mononuclear leukocytes | antigen presenting cells | cerebral cortex cells |
| neuroblastoma cell lines | helper T lymphocytes | cytotoxic T cells |
| dermal cells | sarcoma cell lines | liver cells |
| bone marrow-derived macrophages | plasma cells | fibroblasts |
| peripheral blood monocytes | central nervous system cells | endothelial cells |
| PBMCs | natural killer cells | effector T lymphocytes |
| macrophage cancer cell lines | neurons | naive lymphocytes |
| fibroblast cell lines | thymocytes | astrocytes |
| brain cells | neural stem cells | monocyte-derived dendritic cells |
| keratinocytes | vascular smooth muscle cells | carcinoma cell lines |
| gonadal cells | epidermal cells | osteoclasts |
| B-lymphocyte derived cell lines | monocyte-derived macrophages | smooth muscle cells |
| peripheral blood leukocytes | connective tissue cells | hematopoietic cells |
| memory T lymphocytes | eosinophils | leukemia cells |
| leukocyte cell lines | neutrophils | tumor cells |
| epithelial cells | leukemia cell lines | T lymphocytes |
| embryonic cells | Th2 cells | macrophages |
| neuroglia | hematopoietic progenitor cells | cancer cells |
| mast cells | stem cells | muscle cells |
| nervous tissue cell lines | Schwann cells | epithelial cell lines |
| embryonic cell lines | breast cell lines | phagocytes |
| CD4+ T-lymphocytes | heart cells |  |
| neuroblasts | regulatory T lymphocytes |  |
| Th1 cells | tumor cell lines |  |

**Table S2.**
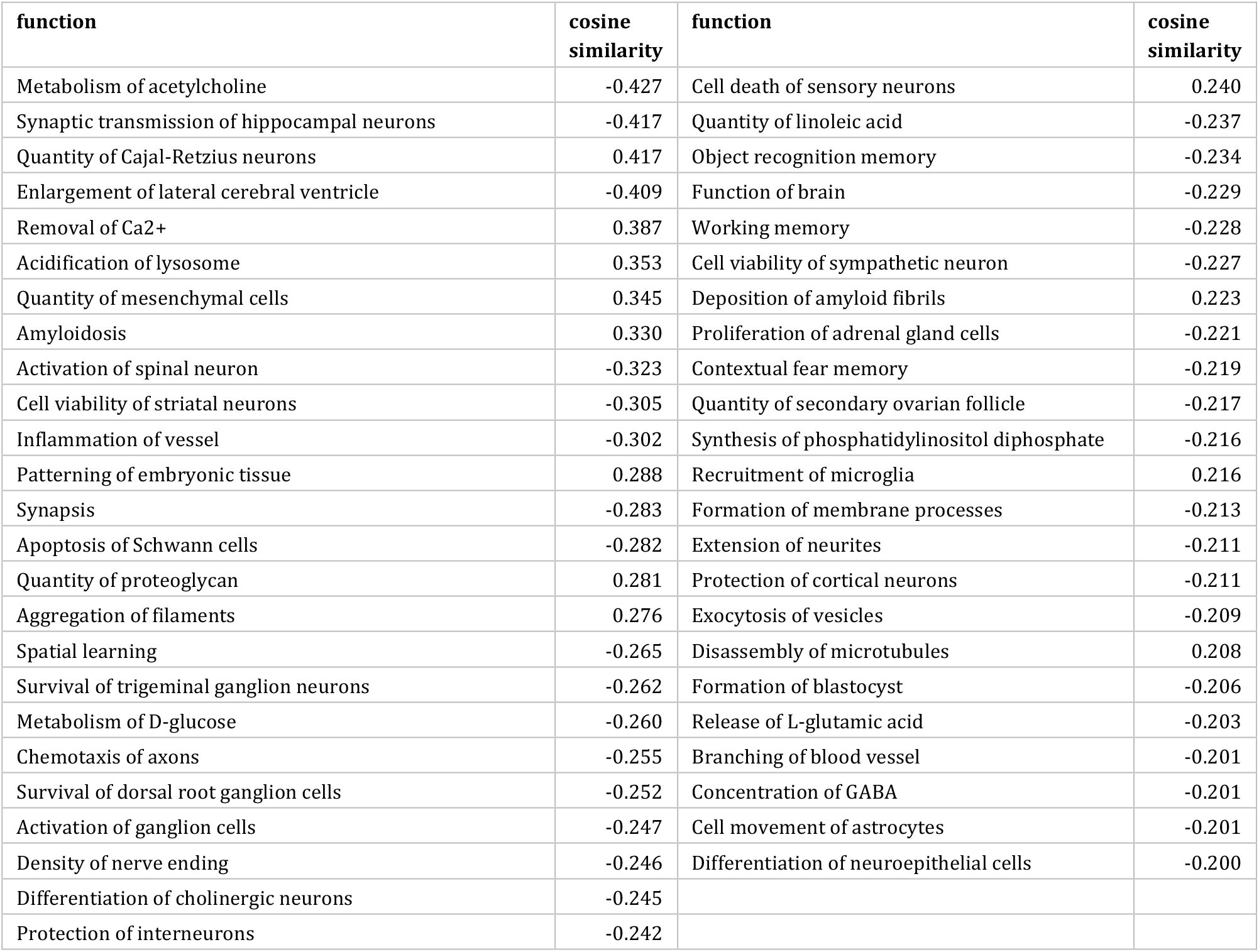
Alzheimer’s disease: top-scoring functions.

| function | cosine similarity | function | cosine similarity |
| --- | --- | --- | --- |
| Metabolism of acetylcholine | -0.427 | Cell death of sensory neurons | 0.240 |
| Synaptic transmission of hippocampal neurons | -0.417 | Quantity of linoleic acid | -0.237 |
| Quantity of Cajal-Retzius neurons | 0.417 | Object recognition memory | -0.234 |
| Enlargement of lateral cerebral ventricle | -0.409 | Function of brain | -0.229 |
| Removal of Ca<sup>2+</sup> | 0.387 | Working memory | -0.228 |
| Acidification of lysosome | 0.353 | Cell viability of sympathetic neuron | -0.227 |
| Quantity of mesenchymal cells | 0.345 | Deposition of amyloid fibrils | 0.223 |
| Amyloidosis | 0.330 | Proliferation of adrenal gland cells | -0.221 |
| Activation of spinal neuron | -0.323 | Contextual fear memory | -0.219 |
| Cell viability of striatal neurons | -0.305 | Quantity of secondary ovarian follicle | -0.217 |
| Inflammation of vessel | -0.302 | Synthesis of phosphatidylinositol diphosphate | -0.216 |
| Patterning of embryonic tissue | 0.288 | Recruitment of microglia | 0.216 |
| Synapsis | -0.283 | Formation of membrane processes | -0.213 |
| Apoptosis of Schwann cells | -0.282 | Extension of neurites | -0.211 |
| Quantity of proteoglycan | 0.281 | Protection of cortical neurons | -0.211 |
| Aggregation of filaments | 0.276 | Exocytosis of vesicles | -0.209 |
| Spatial learning | -0.265 | Disassembly of microtubules | 0.208 |
| Survival of trigeminal ganglion neurons | -0.262 | Formation of blastocyst | -0.206 |
| Metabolism of D-glucose | -0.260 | Release of L-glutamic acid | -0.203 |
| Chemotaxis of axons | -0.255 | Branching of blood vessel | -0.201 |
| Survival of dorsal root ganglion cells | -0.252 | Concentration of GABA | -0.201 |
| Activation of ganglion cells | -0.247 | Cell movement of astrocytes | -0.201 |
| Density of nerve ending | -0.246 | Differentiation of neuroepithelial cells | -0.200 |
| Differentiation of cholinergic neurons | -0.245 |  |  |
| Protection of interneurons | -0.242 |  |  |

**Table S3.** Alzheimer’s disease: top-scoring genes.

| gene | z-score | gene | z-score | gene | z-score | gene | z-score |
| --- | --- | --- | --- | --- | --- | --- | --- |
| PSEN1* | 13.970 | SLC17A6 | 4.459 | YOD1 | 3.722 | HTR1A | 3.291 |
| PSEN2* | 13.545 | NREP | 4.438 | WNT2B | 3.674 | CLASP2 | -3.279 |
| NGF* | -9.819 | CFI | -4.417 | FBXO2 | 3.673 | NTF4 | -3.268 |
| Slfn1 | -7.192 | ALB* | -4.387 | EIF4E2 | 3.648 | Atg5 | 3.225 |
| APOE* | 7.096 | NEUROG2 | -4.274 | IGSF1 | 3.641 | ZNF76 | -3.223 |
| PRR5 | 7.023 | CASP2 | -4.161 | REST | 3.615 | ICMT | -3.219 |
| APP* | 6.482 | MARCHF5 | 4.063 | DAPK3 | 3.597 | AIPL1 | -3.211 |
| INS* | -6.423 | PLOD1 | 4.055 | CDH5 | -3.555 | IRF8 | -3.201 |
| HMGCR* | 6.390 | DLX5 | -4.052 | MAML2 | 3.441 | CYP11B2 | -3.198 |
| BDNF* | -6.165 | INTS11 | -4.051 | MAML3 | 3.441 | LRRN1 | 3.188 |
| CUX2 | 5.992 | SRSF10 | 4.014 | GH2 | 3.434 | AGT | -3.182 |
| SLITRK5 | -5.611 | DLX6 | -4.012 | ZSCAN21 | 3.431 | PDCD5 | -3.176 |
| LOX | 5.509 | PHF5A | -3.987 | CREM | -3.423 | GUCY2F | -3.175 |
| FBXL7 | -5.342 | CPEB3 | -3.972 | PAQR3 | 3.422 | EIF2B2 | -3.172 |
| HRG | -5.089 | GAB1 | -3.948 | IFT20 | 3.402 | PTGIS | 3.165 |
| BAIAP2 | 5.019 | SIM1 | -3.920 | HCK | -3.390 | LLGL2 | 3.158 |
| BID | -4.931 | APPL1 | -3.886 | ADAM19 | -3.386 | PIM1 | -3.133 |
| PSENEN* | 4.894 | TTYH1 | 3.842 | NFIC | -3.378 | CFB | -3.125 |
| HSPE1 | -4.824 | SLC8A1 | -3.823 | MIB1 | 3.371 | BAX | -3.111 |
| SLC30A3 | -4.670 | SV2A | -3.811 | HSP90AA1 | 3.363 | RELN | 3.081 |
| Csl | 4.592 | NEURL1 | -3.802 | ATP2A1 | -3.348 | ALOX12 | -3.069 |
| B4GALNT1 | 4.518 | CST7 | -3.790 | PLAU* | -3.342 | FZD8 | 3.068 |
| AMOT | -4.484 | ARNT2 | -3.760 | RHBDF1 | -3.327 | WNT16 | 3.062 |
| AMOTL2 | -4.484 | BATF2 | -3.756 | SCGB3A2 | 3.326 | PLEKHM1 | 3.059 |
| AMOTL1 | -4.484 | HLA-B | -3.745 | ATF6 | -3.305 | NSF | -3.048 |

Genes marked with (\*) are associated with Alzheimer's disease in the QKB. Genes not marked with (\*) are predicted to be causally associated.

**Table S4.** Pulmonary hypertension: top-scoring functions.

| function | cosine similarity | function | cosine similarity |
| --- | --- | --- | --- |
| Recruitment of eosinophils | 0.337 | Proliferation of effector memory T lymphocytes | 0.210 |
| Muscularization of artery | 0.312 | Differentiation of naive lymphocytes | 0.209 |
| Proliferation of cytotoxic T cells | 0.293 | Binding of stromal cells | 0.208 |
| Secretion of mucus | 0.285 | Synthesis of leukotriene C4 | 0.207 |
| Systolic pressure of right ventricle | 0.284 | Cell viability of monocyte-derived dendritic cells | 0.202 |
| Function of helper T lymphocytes | 0.272 | Remodeling of vascular tissue | 0.201 |
| Transmigration of macrophages | 0.268 | Synthesis of prostaglandin | 0.201 |
| Proliferation of invariant natural killer T cells | 0.264 |  |  |
| Th2 immune response | 0.257 |  |  |
| Generation of bone marrow-derived dendritic cells | 0.252 |  |  |
| Stimulation of mast cells | 0.252 |  |  |
| Activation of natural killer T lymphocytes | 0.250 |  |  |
| Pulmonary fibrosis or aplastic anemia | 0.247 |  |  |
| Quantity of adenosine | 0.246 |  |  |
| Quantity of Th2 cells | 0.245 |  |  |
| Expansion of antigen presenting cells | 0.240 |  |  |
| Cell movement of basophils | 0.236 |  |  |
| Proliferation of Th1 cells | 0.233 |  |  |
| NK cell proliferation | 0.233 |  |  |
| Production of superoxide | 0.223 |  |  |
| Quantity of phagocytes | 0.223 |  |  |
| Development of follicular T helper cells | 0.219 |  |  |
| Maturation of natural killer cells | 0.218 |  |  |
| Effector phase | 0.217 |  |  |
| Stimulation of monocytes | 0.216 |  |  |

**Table S5.** Pulmonary hypertension: top-scoring genes.

| gene | z-score | gene | z-score | gene | z-score | gene | z-score |
| --- | --- | --- | --- | --- | --- | --- | --- |
| ADORA2B* | 9.536 | PAK2 | 4.334 | CEBPZ | 3.620 | PLAT* | -3.293 |
| ADA* | -7.493 | ZNF668 | -4.288 | NAGLU | 3.599 | NCR3LG1 | 3.283 |
| IL18* | 7.135 | NOS3* | -4.204 | NT5E | -3.577 | SMAD5 | -3.271 |
| PTGDR2* | 6.693 | LOC290071 | -4.184 | Havcr1 | -3.551 | CD82 | 3.244 |
| ACVR2A | -6.193 | JAG2 | -4.143 | IL3 | 3.550 | CD180 | 3.237 |
| AKNA | 6.056 | VIP* | -4.082 | HSPB2 | -3.547 | ACVR1 | -3.237 |
| BMPR2* | -6.043 | PTGIR | -4.064 | MOV10L1 | 3.542 | MXD3 | -3.231 |
| RPTOR* | 5.773 | HJV | -4.024 | MS4A1 | -3.532 | EPAS1* | 3.231 |
| APOE* | -4.927 | CSF1 | 4.017 | ADORA1 | -3.492 | NAA30 | -3.205 |
| CCR3 | 4.846 | CAV1* | -3.999 | GRB10 | -3.489 | IL33 | 3.200 |
| AQP11 | -4.839 | CD83 | -3.967 | IL17RB | 3.483 | STOX1 | -3.194 |
| IL15 | 4.838 | POU2AF1 | 3.951 | IFITM3 | -3.474 | TRAF4 | 3.193 |
| ARG1* | 4.813 | CCR6 | 3.930 | ALPL | -3.474 | ANXA13 | -3.178 |
| P4HTM | -4.765 | BID | 3.918 | IL5 | 3.413 | SRFBP1 | -3.171 |
| IL1RN* | -4.741 | CHRNA1 | -3.912 | TMPRSS6 | 3.410 | SETDB1 | 3.165 |
| CSF2* | 4.724 | FBXO11 | -3.905 | Til1 | 3.384 | P2RY1 | -3.160 |
| EGLN1* | -4.678 | BIRC5* | 3.896 | DAZAP2 | -3.373 | TNFRSF4 | 3.119 |
| TNFSF4* | 4.636 | ARNT | 3.863 | BLVRA | -3.352 | PRMT7 | -3.100 |
| KDR* | -4.514 | GDF2* | -3.856 | CD48 | 3.351 | SPI1 | 3.069 |
| CXCR4* | 4.494 | TNFSF10* | 3.818 | IL2RA | -3.339 | CD2 | 3.050 |
| KIF17 | -4.481 | ENDOG | -3.765 | ICOS | 3.331 | TFR2 | -3.045 |
| GREB1 | 4.444 | Pln | -3.706 | WNT2 | 3.324 | CFB | 3.036 |
| ZNF613 | 4.441 | Adora3/LOC100911796 | 3.676 | IL9R | 3.315 | IRF8 | 3.020 |
| NOS2* | 4.438 | CAMTA1 | 3.676 | STAT1 | 3.305 | HIF1AN | -3.016 |
| CLSTN1 | 4.391 | ZNF260 | 3.676 | FOSL2* | 3.298 | USP22 | 3.013 |

Genes marked with (\*) are associated with pulmonary hypertension in the QKB. Genes not marked with (\*) are predicted to be causally associated.

**Table S6.** Psoriasis: top-scoring functions.

| function | cosine similarity | function | cosine similarity | function | cosine similarity | function | cosine similarity |
| --- | --- | --- | --- | --- | --- | --- | --- |
| Influx of neutrophils | 0.414 | Activation of synovial fibroblasts | 0.306 | Accumulation of pyruvic acid | -0.275 | Proliferation of colony-forming granulocyte-macrophages | 0.253 |
| Activation of Th17 cells | 0.394 | Adhesion of mesenchymal stem cells | 0.304 | Necroptosis of bone marrow-derived macrophages | 0.274 | I-kappaB kinase/NF-kappaB cascade | 0.253 |
| Migration of Langerhans cells | 0.393 | Synthesis of 5,6,7,8-tetrahydrobiopterin | 0.303 | Permeability of tight junctions | 0.273 | Neurogenesis of neural stem cells | 0.253 |
| Th17 immune response | 0.386 | Fever | 0.302 | Activation of mesangial cells | 0.272 | Proliferation of fibroblast-like synoviocytes | 0.253 |
| Quantity of IL-1a in blood | 0.371 | Immune response of brain | 0.302 | Release of L-cysteine | 0.272 | Generation of Th9 cells | -0.253 |
| Activation of Th1 cells | 0.370 | Killing of Haemophilus influenzae | 0.295 | Inflammation of absolute anatomical region | 0.272 | Binding of Sertoli cells | 0.252 |
| Loss of proteoglycan | 0.368 | Killing of Listeria monocytogenes 10403S | 0.295 | Binding of E. coli | 0.271 | Trafficking of B lymphocytes | 0.252 |
| Production of anti-DNA antibody | 0.357 | Polarization of T-cell hybrid cells | 0.295 | Clearance of Pseudomonas aeruginosa | 0.270 | Binding of microvessel | 0.251 |
| Quantity of nitric oxide | 0.354 | Release of prostaglandin E2 | 0.293 | Growth of Mycobacterium tuberculosis | -0.269 | Acute phase reaction | 0.251 |
| Formation of nitrite | 0.353 | Proinflammatory response | 0.293 | Dissemination of Klebsiella pneumoniae | -0.269 | Efflux of sphingomyelin | -0.250 |
| Apoptosis of thyroid cells | 0.337 | Fragmentation of DNA fragment | 0.292 | Formation of PML nuclear bodies | 0.269 | Activation of alveolar macrophages | 0.250 |
| Synthesis of leukotriene B4 | 0.336 | Clearance of Staphylococcus aureus | 0.292 | Activation of myeloid-derived suppressor cells | 0.268 | Induction of muscle cells | 0.250 |
| Translocation of granules | 0.335 | Cellular infiltration by CD4+ T-lymphocytes | 0.292 | Induction of follicular T helper cells | 0.267 | Increased localization of AST | 0.249 |
| Stimulation of lung cells | 0.335 | Stimulation of hepatocytes | 0.287 | Activation of bone marrow-derived dendritic cells | 0.266 | Quantity of enterobacteriaceae | -0.249 |
| Cell movement of naive B cells | 0.334 | Apoptosis of microglia | 0.286 | Stimulation of chondrocytes | 0.264 | Apoptosis of trophoblast cells | 0.248 |
| Adhesion of peripheral blood monocytes | 0.332 | Proliferation of endometriotic stromal cells | 0.285 | Damage of genitourinary system | 0.264 | Outgrowth of lymph vessel | 0.247 |
| Degradation of connective tissue | 0.323 | Release of sphingolipid | 0.284 | Release of non-esterified fatty acid | 0.263 | Apoptosis of effector memory T lymphocytes | 0.247 |
| Stimulation of airway smooth muscle cells | 0.319 | Response of skeletal muscle | 0.282 | Induction of histamine | 0.263 | Chemotaxis of Th2 cells | 0.246 |
| Scaling of skin | 0.317 | Permeability of microvasculature | 0.280 | Activation of sensory neurons | 0.261 | Maturation of myeloid dendritic cells | 0.246 |
| Activation of keratinocytes | 0.316 | Chemotaxis of basophils | 0.280 | Release of cyclic GMP | 0.260 | Binding of C/ebp beta binding site | 0.243 |
| Induction of prostaglandin | 0.312 | Quantity of apoptotic endocrine cell lines | 0.280 | Writhing | 0.256 | Apoptosis of islets of Langerhans | 0.243 |
| Cell movement of memory B cells | 0.310 | Response of fibroblasts | 0.279 | Survival of Francisella tularensis | -0.255 | Stimulation of vascular endothelial cells | 0.242 |
| Concentration of epoprostenol | 0.310 | Hematopoiesis of myeloid progenitor cells | -0.279 | Activation of natural killer T lymphocytes | 0.255 | Apoptosis of placenta | 0.241 |
| Inflammatory infiltrate | 0.307 | Release of reactive oxygen species | 0.278 | Killing of Leishmania | 0.254 | Chemoattraction of eosinophils | 0.241 |
| Cell viability of oligodendrocyte precursor cells | -0.307 | Stimulation of hyaluronic acid | 0.277 | Binding of vascular smooth muscle cells | 0.253 | Flux of Ca<sup>2+</sup> | 0.240 |

**Table S7.** Psoriasis: top-scoring genes.

| gene | z-score | gene | z-score | gene | z-score | gene | z-score |
| --- | --- | --- | --- | --- | --- | --- | --- |
| KLK5 | 9.014 | GABRA2 | 5.358 | FILIP1 | -3.917 | STAT1 | 3.634 |
| DTL* | -8.339 | ACKR2* | -5.264 | SIGIRR* | -3.891 | CCL20* | 3.609 |
| IL36A* | 8.312 | IL1B | 5.150 | ARRB1 | 3.884 | S100A4 | 3.607 |
| IL17C* | 8.289 | IL10* | -5.130 | TGFB1* | 3.880 | AGER | 3.603 |
| TEK* | 7.871 | IL22RA2* | -5.038 | IL12RB2 | -3.862 | IFNL1 | 3.564 |
| BANF1 | -7.736 | IL22* | 4.824 | PRKCA | 3.831 | CCR2 | 3.558 |
| NFKBIZ | 7.492 | TCL1A | 4.761 | PLAUR | 3.823 | TNFSF12 | 3.547 |
| PLA2G2D* | -7.490 | IL4* | -4.713 | S100A8 | 3.822 | IFNL3 | 3.513 |
| IL17A* | 7.329 | CNPY3 | 4.676 | Tcrd | 3.818 | CYFIP2 | -3.500 |
| IL23A* | 7.180 | CXCL8* | 4.535 | Rcan1 | 3.818 | NPC1 | -3.493 |
| IL1RL2 | 6.931 | CFP | 4.534 | PNPT1 | -3.789 | CD200 | -3.486 |
| TNF* | 6.862 | IRF7 | 4.482 | IL1A | 3.771 | IL12B | 3.481 |
| TNIP1 | -6.362 | CXCR5 | 4.471 | IL20* | 3.754 | PF4 | 3.470 |
| HSD17B14 | -6.263 | SCGB1A1 | -4.411 | SWAP70 | 3.751 | TNFAIP3 | -3.445 |
| IL1RN* | -6.240 | NPS | 4.385 | IL1RL1 | -3.740 | TRHR | 3.445 |
| ADAMTS12 | -5.942 | Wfdc17 | -4.383 | CSF3R | -3.738 | GLIS1 | 3.415 |
| CXCL5* | 5.931 | TRAIP | -4.364 | IL36G | 3.726 | HSP90B1 | 3.414 |
| CXCL17 | -5.741 | ZFP36 | -4.339 | CXCL1* | 3.718 | RBBP4 | 3.402 |
| Ccl2* | 5.738 | TRU-TCA1-1 | -4.261 | OPA1 | -3.712 | Usp17la | 3.392 |
| CAMP | 5.652 | CCR1 | 4.205 | C7 | 3.669 | RGS10 | -3.375 |
| KDR | -5.582 | IRF3 | 4.113 | STAT6 | -3.663 | ZBTB46 | -3.373 |
| GPR34 | 5.424 | PTPN22 | -4.100 | PTPRT | -3.653 | TRAF3IP2* | 3.372 |
| IFNG | 5.396 | CEBPE | 4.048 | Saa3 | 3.650 | IL23R | 3.333 |
| Ctf2 | 5.393 | CXCL2 | 3.965 | APOA1 | -3.646 | ANKRD17 | 3.330 |
| REG3A | 5.393 | MFAP2 | -3.941 | APP | 3.643 | PGRMC1 | 3.318 |

Genes marked with (\*) are associated with psoriasis in the QKB. Genes not marked with (\*) are predicted to be causally associated.

